# DAPE-PD: Diffusion adaptation of genomic foundation model enables tailored-generation of *cis*-regulatory DNA

**DOI:** 10.64898/2026.10.24.761598

**Authors:** Martin Echavarria Galindo, Shansan Gong, Yuewei Zhan, Shuchen Wang, Lingpeng Kong, Yong Lai

## Abstract

Genomic foundation models (GFMs) have transformed our ability to learn the evolutionary language encoded in genomes, yet their potential for programmable regulatory DNA engineering remains underexplored. Regulatory function depends on combinatorial interactions among distributed sequence elements, motivating generation with bidirectional context. Here, we introduce DAPE-PD (<u>D</u>iffusion <u>A</u>daptation of <u>P</u>retrained <u>E</u>vo for <u>P</u>romoter <u>D</u>esign), which adapts the autoregressive Evo model into a bidirectional masked diffusion model without foundation-scale retraining. DAPE-PD combines pretrained genomic representations with iterative unmasking to generate regulatory DNA sequences. Following genus-specific fine-tuning on genome-derived intergenic sequences, DAPE-PD generated sequences that retained genus-associated compositional features despite low similarity to the fine-tuning datasets. Experimental validation identified functional de novo promoters, while in silico prioritization recovered active designs from small experimental candidate sets across three non-model gut commensals. This framework connects genomic pretraining with diffusion-based generation, extending GFMs from sequence understanding toward functional *cis*-regulatory DNA engineering in living cells.

## Introduction

Engineering microbial functions requires regulatory elements that operate predictably in the intended host. However, genetic toolkits and experimentally characterized regulatory parts are not available for many of the abundant commensal bacteria in the human gut microbiome^1,2^. Promoters not only determine the strength, stability, and timing of engineered gene expression, but in the highly fluctuating intestinal environment their compatibility with the host transcriptional machinery ultimately determines whether a genetic circuit functions at all^3,4^. Existing strategies based on motif-guided rational design^5–7^ or random mutagenesis and library screening^4,5,8,9^ have produced regulatory parts that typically require extensive experimentation; such strategies are difficult to apply to organisms for which functional regulatory data is limited. The lack of programmable *cis*-regulatory elements for non-model bacteria with key roles in human health therefore remains a fundamental bottleneck to genetic circuit design and microbiome engineering.

Recent advances in deep learning have created new opportunities for regulatory sequence engineering. Attention-based neural architectures provide expressive sequence representations and improve predictive accuracy for genomic tasks^10–12^. These attention-based models have been used to decipher regulatory DNA logic, and genomic language models (gLMs) are being trained across vast sequence datasets to infer the underlying "grammar" and motif conservation patterns of DNA^13,14^.

gLMs have adopted next-token prediction objectives similar to those developed for autoregressive (AR), decoder-only architectures in large language models (LLMs) to generate DNA sequences de novo. Evo 1, a 7-billion-parameter AR model trained on approximately 300 billion nucleotides from 80,000 prokaryotic genomes, is the most comprehensive prokaryotic gLM to date^15^. It features single-nucleotide tokenization and enables the zero-shot prediction of mutation effects and expression via log-likelihood scoring. Evo 1 has also been used to design functional systems (e.g., operons and gene clusters)^16^ and whole bacteriophage genomes^17^.

Although these studies have established large-scale genomic pretraining as a source of biological prior knowledge, AR gLMs are architecturally constrained to left-to-right unidirectional sequence processing and generation. The use of a causal attention mask during training prevents the leakage of future token information but also prevents global context capture (i.e. left-to-right and right-to-left information) of the input sequences. This unidirectional paradigm remains fundamentally mismatched with the bidirectional, flanking regulatory dependencies that govern genomic DNA. In biological systems, promoter function involves coordinated interactions among sequence features distributed across an entire regulatory region, including recognition motifs, spacing, local composition and higher-order context, rather than from a strictly sequential process^18,19^.

Diffusion models, alternatively, leverage bidirectional denoising to reconstruct signals from a noisy distribution using a global sequence context. The generative performance and output diversity of such models thus surpass those of AR approaches^20^. Recent studies have begun to explore diffusion models for biological sequence generation^21,22^. However, current diffusion-based DNA models lack the scale and phylogenetic diversity of Evo 1, restricting their biological scope.

Training diffusion models from scratch at scale requires enormous computational resources, limiting their accessibility and practical deployment. To overcome this limitation, we recently developed a general framework for adapting pretrained AR language models to work as diffusion language models through efficient fine-tuning^23^. Integrating large-scale AR pretraining with diffusion-based generation offers a previously unexplored avenue to repurpose these models for programmable regulatory DNA engineering.

Here, building upon this adaptation framework, we developed DAPE-PD (<u>D</u>iffusion <u>A</u>daptation of <u>P</u>retrained <u>E</u>vo for <u>P</u>romoter <u>D</u>esign), which represents the first diffusion-based adaptation of a genomic foundation model. DAPE-PD converts the pretrained Evo 1 from an AR model to a bidirectional masked diffusion model. DAPE-PD generated high-quality de novo promoters for model organisms and gut-associated commensals. These promoters display substantially greater sequence novelty than those generated by conventional AR fine-tuning, while retaining native-like compositional features. Coupled with computational prioritization, the framework recovered active promoters from small experimental candidate sets in data-scarce non-model bacteria. These results establish a practical route from genomic foundation models to programmable *cis*-regulatory DNA engineering without requiring foundation-scale retraining or large-scale experimental validation.

## Results

### Diffusion adaptation of Evo 1 for de novo promoter design

Conventional deep learning approaches for de novo promoter design have largely depended on well-curated datasets from model organisms, limiting their applicability to non-model species^24,25^. To overcome this data scarcity, we constructed a specialized training set comprising upstream regulatory sequences (50–200 bp) extracted from complete genomes of target genera, enabling gLM fine-tuning where large-scale data are unavailable (**Fig. 1a**, Methods). Fine-tuning these genus-specific intergenic sequences enabled the model to capture dataset-specific regulatory features while retaining Evo 1’s general representational capacity.

**Fig. 1.**
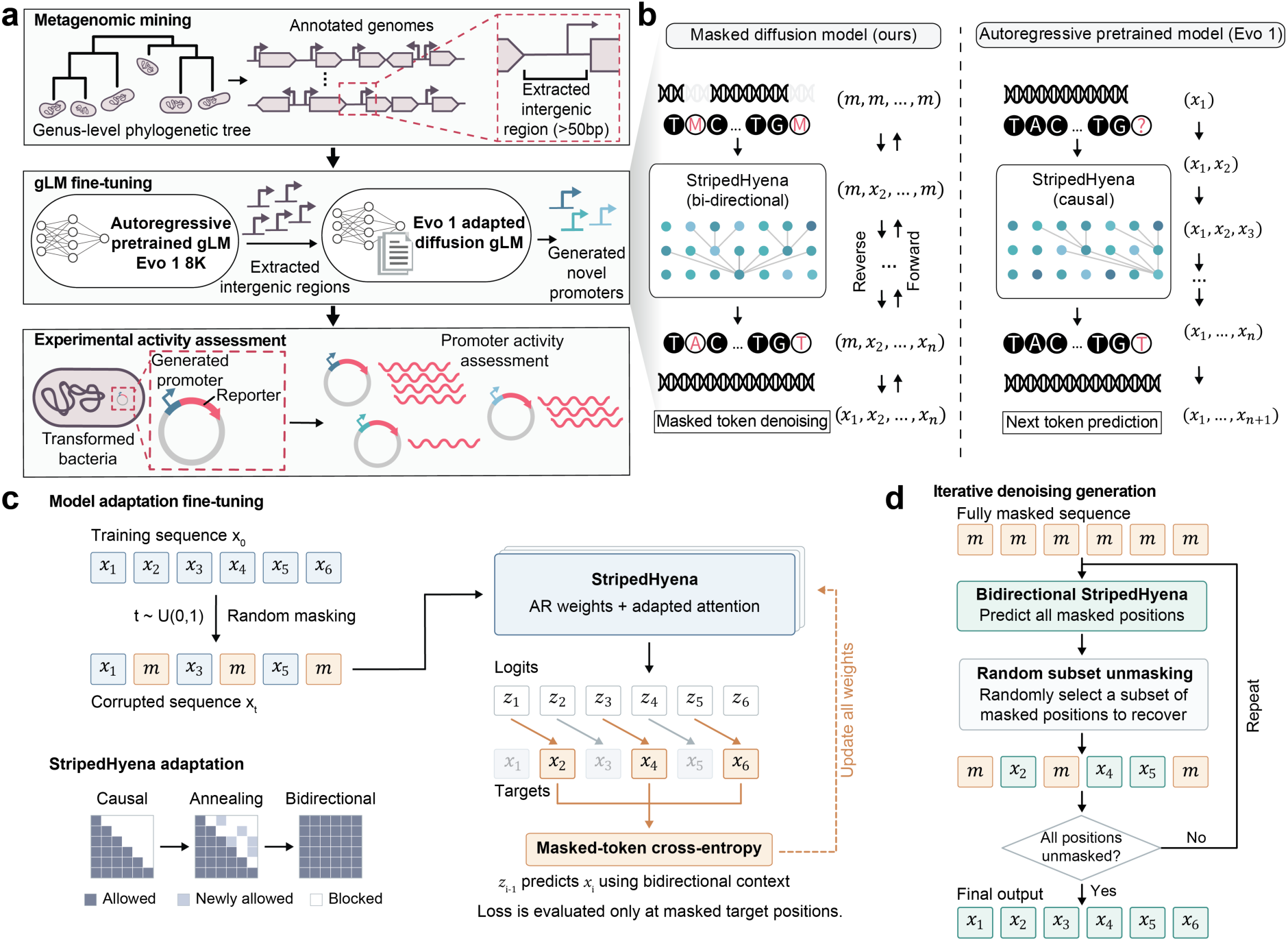
A diffusion-adapted genomic foundation model pipeline for the generation of functional de novo promoters. **a**, Overview of integrated pipeline: from genus-level intergenic sequence mining and gLM fine-tuning to promoter generation and experimental validation. **b**, Comparison of masked diffusion and AR generation. Unlike unidirectional AR models, the adapted diffusion model leverages the bidirectional context for de novo generation (ATCG, standard DNA bases; *m*, masked positions; *x*, specified nucleotides). **c**, Overview of our approach for adapting the autoregressive Evo 1 model into a bidirectional masked diffusion model. During fine-tuning, the causal mask is gradually removed, ultimately rendering the model bidirectional as it leverages bidirectional context to predict each masked token. Training sequences *x*_0_ are corrupted to *x_t_* by masking tokens via a forward process governed by the diffusion timestep *t*. The shift operation on the output logits is defined such that the logits at each position are aligned with the prediction target of the next token, i.e. *z_i_*_−1_ is used to predict *x_i_*. Orange connections identify masked targets contributing to the cross-entropy loss; dashed arrows indicate parameter updates. Attention matrices depict permitted query-key access. **d**, Generation begins with a fully masked sequence that is progressively denoised, with each step randomly recovering a subset of predicted positions while the remainder stay masked. Blue, orange, and teal token boxes denote observed, masked and generated nucleotides, respectively.

During fine-tuning, we converted the pretrained AR Evo 1 model into a masked diffusion model by replacing its causal attention mechanism with bidirectional attention (**Fig. 1b,c**). During training, a subset of input tokens was randomly masked (denoted by "M" in **Fig. 1b**), and the model was optimized to reconstruct the original sequences via cross-entropy loss. Gradients from this loss were used to update the model parameters (**Fig. 1c**).

Generation proceeded in a fully diffusion-based manner, starting from sequences composed entirely of masked tokens (**Fig. 1d**). At each iteration, the model predicted all masked positions, randomly recovered a subset, and retained the rest as masked, continuing until no masked tokens remained. This iterative denoising produced sequences that were both novel and syntactically consistent with the genomic “grammar” of the fine-tuning datasets.

### Diffusion increases promoter novelty while preserving host sequence grammar

To benchmark our diffusion-based pipeline against conventional approaches, we first fine-tuned the pretrained Evo 1 model using a next-token prediction objective on three model species for which high-throughput functional data provided validated reference standards^4^: *Escherichia coli* (Gram-), *Pseudomonas aeruginosa* (Gram-) and *Bacillus subtilis* (Gram+). As AR baselines, we compared two AR fine-tuning strategies: joint multi-task training (hereafter referred to as joint AR fine-tuning) across all three species, and separate genus-specific fine-tuning (hereafter referred to as separate AR fine-tuning).

For model bacteria genus-level mining, we retrieved all complete genomes and annotations from NCBI for these three genera, yielding an initial collection of 8,966 *Escherichia* genomes, 4,311 *Pseudomonas*, and 3,630 *Bacillus* (**Fig. 2a**). From these genomes, we extracted 50–200 bp intergenic regions and removed redundancies via Linclust^26^ (**Fig. 2b**), resulting in non-redundant datasets of 366,220 *Escherichia* (mean: 165 bp; **Extended Data Fig. 1a,b**), 1,884,218 *Pseudomonas* (mean: 159 bp; **Extended Data Fig. 1c,d**), and 748,056 *Bacillus* (mean: 163 bp; **Extended Data Fig. 1e,f**) intergenic sequences.

**Fig. 2.**
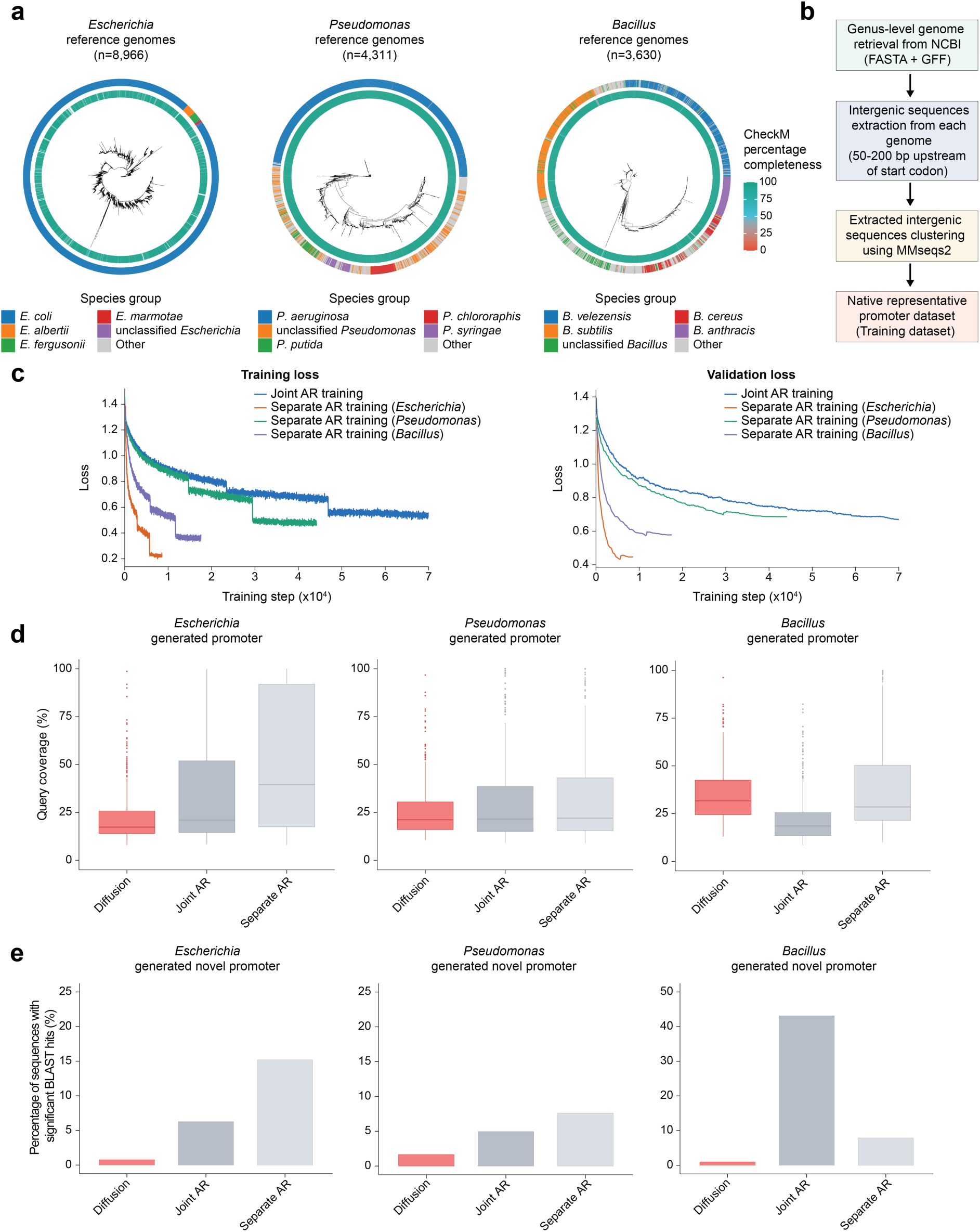
Generative design of promoters in model bacteria. **a**, Inferred phylogenetic tree of *Escherichia*, *Pseudomonas* and *Bacillus* genomes used for promoter mining, annotated with NCBI-reported CheckM completeness scores. **b**, Workflow for mining intergenic sequences at the genus level to construct datasets for model fine-tuning. **c**, Training and validation loss curves for joint versus separate AR fine-tuning. **d**, Mean BLAST query coverage of promoters generated by diffusion, joint AR and separate AR fine-tuning against the corresponding training sequences from three model bacterial genera. **e**, Percentage of low-similarity sequences (<30% query coverage) returning significant BLAST hits (E-value < 0.05) against the full NCBI database.

Joint AR fine-tuning produced consistently higher training and validation losses than the three separate AR fine-tuning experiments (**Fig. 2c**), suggesting that multi-task learning compromises specificity because of inter-species sequence divergence. Analysis of the generated sequences showed that, across all three genera, promoters from joint AR fine-tuning exhibited markedly lower mean query coverage against the training dataset than those from separate AR fine-tuning. Mean query coverage under joint versus separate AR fine-tuning was 47.72% versus 61.20% for *Escherichia* (**Fig. 2d** and **Extended Data Fig. 1g**), 34.80% versus 39.10% for *Pseudomonas* (**Fig. 2d** and **Extended Data Fig. 1h**), and 21.90% versus 49.12% for *Bacillus* (**Fig. 2d** and **Extended Data Fig. 1i**). Despite these differences, both next-token prediction approaches generated a substantial proportion of promoters that were similar to sequences in the training data. To evaluate functional viability, we screened low training dataset similarity sequences (<30% query coverage) for species-specific housekeeping σ^70^ factor binding motifs using FIMO^27^. Motif recovery for joint versus separate AR fine-tuning was 7.05% versus 7.60% in *Escherichia*, 1.32% versus 0.72% in *Pseudomonas*, and notably divergent at 5.16% versus 25.0% in *Bacillus*.

Having established that AR fine-tuning of Evo 1 generates functional promoters, we next tested whether a diffusion-based fine-tuning pipeline could optimize generation quality along two key axes: sequence novelty and functional viability. We therefore adapted Evo 1 via diffusion fine-tuning using the intergenic datasets from the same three genera. Fine-tuning was performed in a genus-specific manner, given that separate AR fine-tuning had obtained a lower loss and outperformed joint training for all three datasets.

Overall, diffusion-based generation yielded promoters with significantly greater novelty than the AR counterpart, as evidenced by lower training dataset mean query coverage in *Escherichia* (23% vs. 61.20% separate AR; **Fig. 2d** and **Extended Data Fig. 1g**), *Pseudomonas* (25.83% vs. 39.10%; **Fig. 2d** and **Extended Data Fig. 1h**), and *Bacillus* (35.2% vs. 49.12%; **Fig. 2d** and **Extended Data Fig. 1i**). Comprehensive BLAST screening of low-similarity sequences (<30% query coverage) against the full NCBI database confirmed this enhanced novelty, with the diffusion model yielding substantially fewer significant hits than joint or separate AR models in *Escherichia* (0.76% vs. 6.27% and 15.2%), *Pseudomonas* (1.66% vs. 4.96% and 7.61%), and *Bacillus* (0.95% vs. 43.15% and 7.89%) (**Fig. 2e**). Thus, diffusion had accessed sequence space that was poorly represented in either the fine-tuning corpus or known genomic sequences. This reduced native homology is vital for designing orthogonal promoters with minimized regulatory cross-talk.

Greater novelty did not come at the expense of native-like composition. Diffusion-generated low-similarity sequences closely matched genus-specific GC-content distributions, with Wasserstein distances of 0.64 for *Escherichia* (3.05 joint and 0.69 separate AR; **Fig. 3a**), 0.42 for *Pseudomonas* (3.26 joint and 1.85 separate AR; **Extended Data Fig. 2a**), and 0.59 for *Bacillus* (10.5 joint and 2.38 separate AR; **Extended Data Fig. 2b**), and reproduced native 6-mer frequencies with strong correlations (**Fig. 3b** and **Extended Data Fig. 2c,d**). Sigma-factor motif analysis provided an independent measure of regulatory plausibility. Among the low-homology sequences generated, expected promoter motifs were recovered in 12.72% of *Escherichia*, 1.65% of *Pseudomonas* and 25.46% of *Bacillus* diffusion designs, exceeding both AR baselines in *Escherichia* and *Pseudomonas* and slightly exceeding the separate AR baseline in *Bacillus*.

**Fig. 3.**
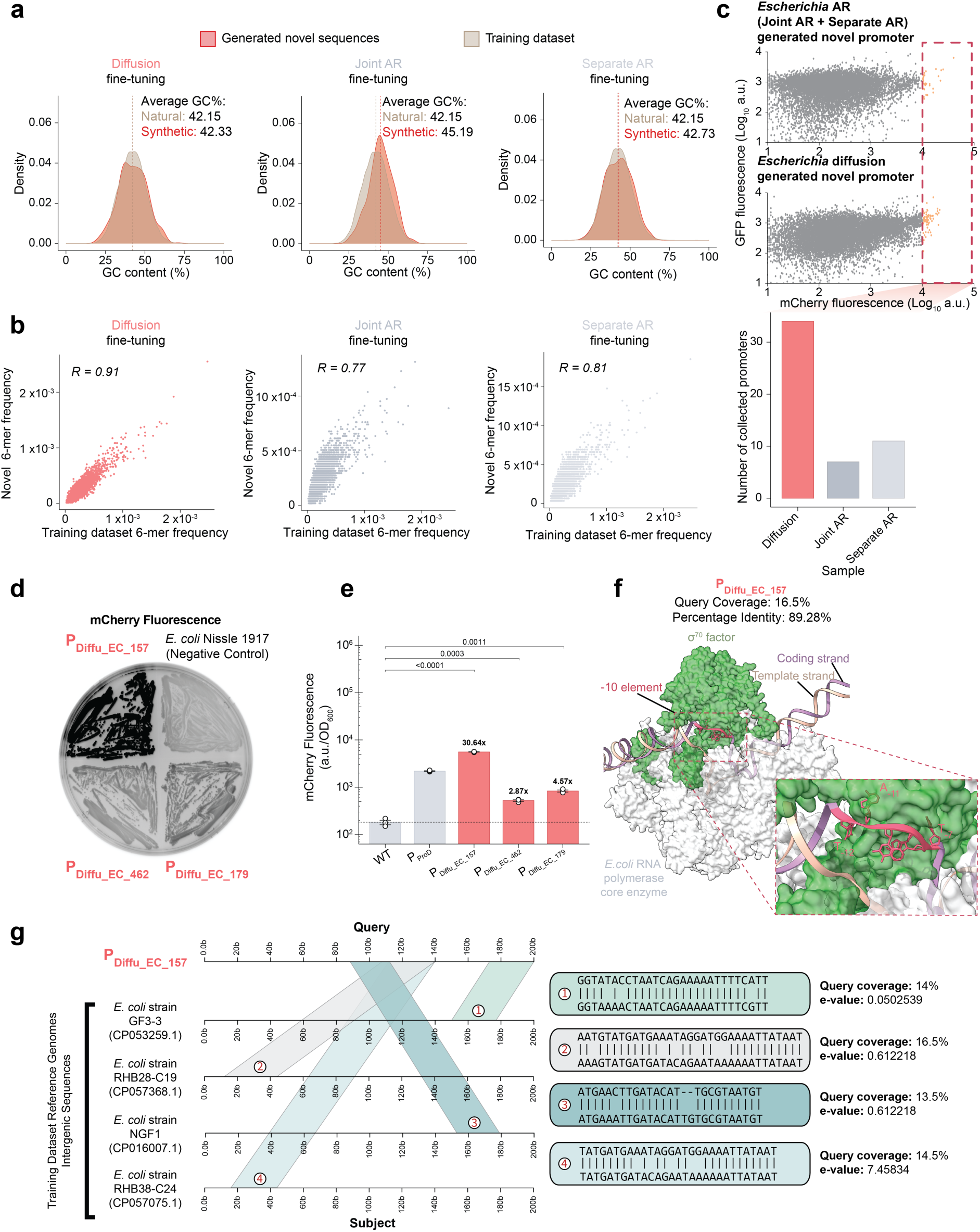
Characterization and functional validation of generated *Escherichia* promoters. **a**, GC-content distributions of generated novel promoters (dotted lines, sample means). **b**, Correlation of 6-mer frequencies between generated novel and native sequences. The effect size is reported as Pearson’s correlation coefficient. **c**, FACS-seq functional enrichment of *Escherichia* promoter libraries in *E. coli* Nissle 1917 (n=10,000 events per sample). Diffusion-generated promoters were significantly enriched (>1000 reads) in the highest mCherry population (orange), reflecting robust transcriptional strength. **d,** mCherry fluorescence levels of three high-performing, diffusion-generated promoters selected based on read counts in *E. coli* Nissle 1917. **e,** Activity profiles of the generated promoters compared to that of the native reference promoter P_ProD_. The fold-change in fluorescence relative to the wild-type (WT) baseline is indicated in bold. Bars show mean ± SE; white circles represent individual biological replicates (n = 3 per group). Statistical significance was determined using an unpaired, two-sided Welch t-test. **f**, Predicted binding model of the *E. coli* RNA polymerase core (grey) and σ⁷⁰ (green) in complex with the top synthetic promoter, PDiffu_EC_157 (-10 element in red). **g,** Representative sequence alignments of PDiffu_EC_157 against the training corpus, highlighting the integration of short regulatory motifs derived from diverse genomes.

We then asked whether these sequence-level differences corresponded to promoter function. Using fluorescence-activated cell sorting (FACS) based pooled screening in *E. coli* Nissle 1917, we quantified the number of promoters from each generation strategy that were enriched in the highest-fluorescence population. For experimental screening, candidates were grouped into two synthesized libraries: a dedicated diffusion-generated library and a combined AR library pooling both separate and joint AR designs. To enable a fair comparison across strategies, we focused our analysis on an equal number (n = 100) of low-homology candidates from each library (see Methods) selected using Promoter Calculator^28^ (see Methods). Under this equalized comparison, diffusion-generated promoters were strongly enriched in the high-fluorescence population: 34 diffusion-generated promoters were recovered, compared with 11 from separate and 7 from joint AR fine-tuning (**Fig. 3c**).

We validated three diffusion-generated *E. coli* promoters with low training dataset similarity and high FACS-seq read counts. PDiffu_EC_157 (query coverage: 16.5%; sequence identity: 89.286%), ranked second by read count, showed the highest activity, substantially exceeding wild-type baseline (**Fig. 3d,e**). In contrast, two lower-ranked variants exhibited reduced expression: PDiffu_EC_462 (query coverage: 16%; sequence identity: 91.667%) and PDiffu_EC_179 (query coverage: 16.667%; sequence identity: 90.909%).

The low sequence homology of P_Diffu_EC_157_ raised the question of whether the generated sequence could nevertheless support canonical transcription initiation. AlphaFold3^29^ modeling of the promoter with the *E. coli* RNA polymerase holoenzyme predicted a high-confidence complex (ipTM = 0.79, pTM = 0.83; **Fig. 3f**) revealing canonical σ⁷⁰ engagement: the −10 element (T_-12_ATAAT_-7_) interacts with the σ2 domain via flipped A_-11_ and T_-7_ nucleotides, accompanied by a ∼15-nucleotide transcription bubble^30^. Although they do not establish mechanisms, structural predictions combined with functional data are consistent with the model in which short regulatory features are recombined into sequence backgrounds that are globally distinct from natural promoters. Sequence alignment confirmed that small native motifs had been retained, indicating selective learning of core promoter features from diverse intergenic sequences (**Fig. 3g** and **Extended Data Fig. 3a,b**).

### Promoters generated by diffusion for non-model bacteria

Having established that our diffusion pipeline yields functionally superior promoters with low sequence similarity relative to AR fine-tuning, we extended this approach to three species of gut bacteria currently lacking well-characterized genetic parts: *Lactiplantibacillus plantarum* (*L. plantarum*), *Bacteroides thetaiotaomicron* (*B. thetaiotaomicron*), and *Bifidobacterium longum* subspecies *infantis* (*B. infantis*)*. L. plantarum*, a probiotic used in industrial fermentation, contributes to intestinal barrier restoration^31^. *B. thetaiotaomicron*, a highly prevalent human gut commensal, was chosen for its long-term colonization stability and potent anti-inflammatory properties^32^, which make it a good candidate for engineered live biotherapies. *B. infantis* is an early colonizer of the infant gut that stimulates immune development and exerts anti-inflammatory effects by metabolizing human milk oligosaccharides and producing short-chain fatty acids^33^.

We mined complete NCBI genomes and annotations from these three genera (*Bacteroides*, n = 409; *Bifidobacterium*, n = 628; *Lactobacillus*, n = 538; **Fig. 4a**) and supplemented them with genus-specific phages (213, 26, and 156 genomes, respectively; **Extended Data Fig. 4**). Intergenic region clustering yielded non-redundant training datasets of 91,698 *Bacteroides* (mean: 158 bp), 72,142 *Bifidobacterium* (mean: 165 bp), and 54,184 *Lactobacillus* (mean: 158 bp) promoters with which to fine-tune our diffusion model (**Fig. 4b**).

**Fig 4.**
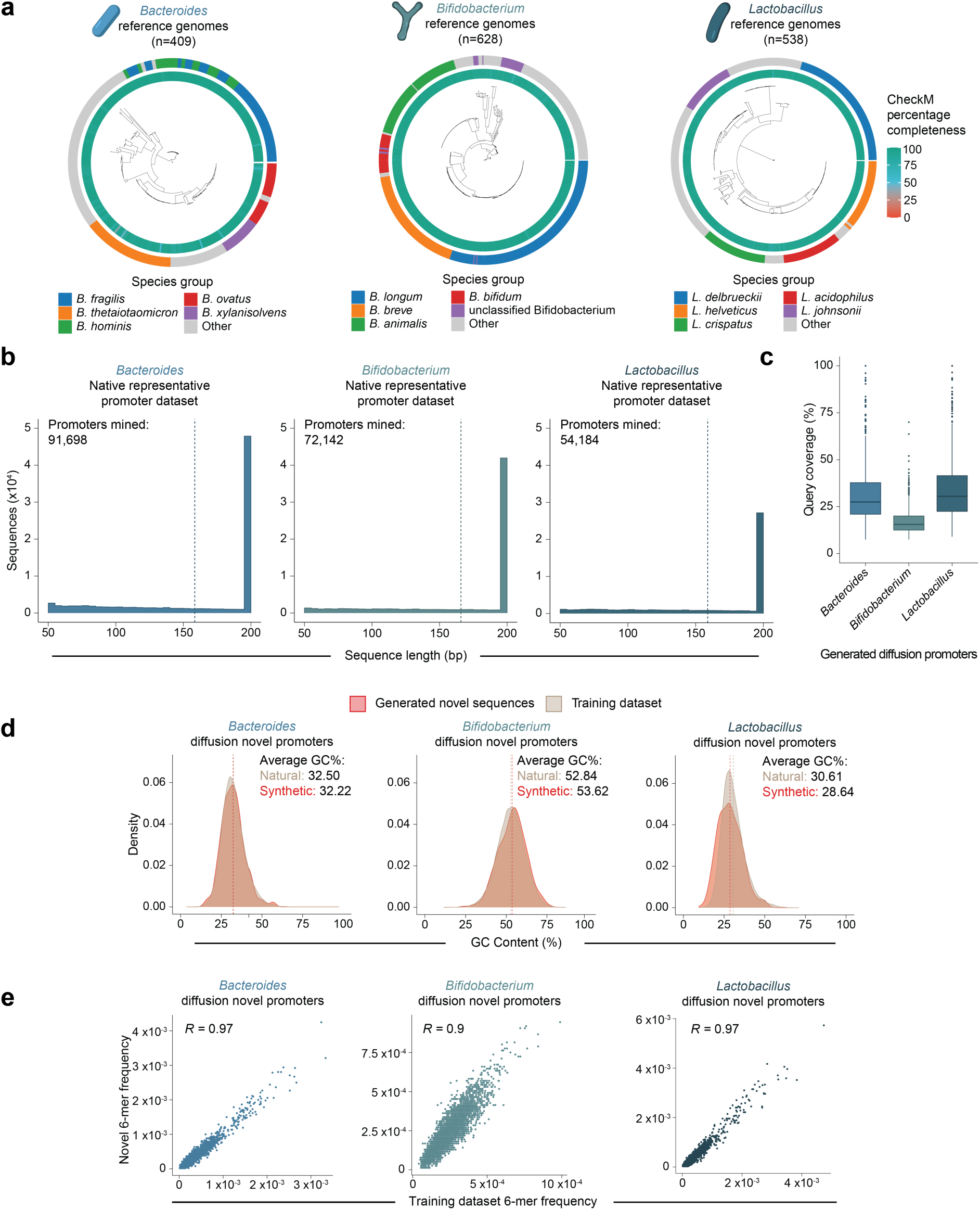
Generative design of diffusion-generated promoters in non-model gut bacteria. **a,** Phylogenetic reference trees for *Bacteroides*, *Bifidobacterium*, and *Lactobacillus* used to source native intergenic sequences, with NCBI-reported CheckM completeness scores and taxonomic annotations. **b,** Length distribution of the curated native *Bacteroides*, *Bifidobacterium*, and *Lactobacillus* promoter dataset used (dotted line, mean). **c,** BLAST query coverage of generated sequences against the corresponding fine-tuning datasets. **d** and **e**, Comparison of native and generated novel sequence composition by GC-content distributions (**d**) and 6-mer frequencies (**e**). Dotted lines indicate sample averages. The effect size is reported as Pearson’s correlation coefficient.

Despite the smaller datasets, the generated sequences retained the same combination of novelty and host-like composition observed in the model genera described above. Mean query coverages against the corresponding fine-tuning sets were 32.42% for *Bacteroides*, 17.82% for *Bifidobacterium*, and 34.71% for *Lactobacillus* (**Fig. 4c**). GC-content distributions and 6-mer frequencies of low-similarity sequences (<30% query coverage) closely tracked those of the native datasets (**Fig. 4d,e**), while only 1.04%, 0.66% and 4.20% of these sequences, respectively, produced significant matches to the full NCBI database. These results confirm that for phylogenetically diverse non-model hosts, our pipeline consistently generates highly novel regulatory elements that preserve native compositional constraints.

Experimental throughput, rather than sequence generation, is often the limiting step for the engineering of non-model organisms^34^. We therefore developed a three-stage prioritization strategy that reduced hundreds of the generated candidates to a small set for testing (**Fig. 5a**). First, to ensure novelty, we retained only those sequences composed entirely of canonical nucleotides (A, C, G, T) that exhibited <30% BLAST query coverage when aligned against the training dataset. Second, using FIMO, we screened for genus-specific regulatory motifs^27^. These reference motifs were empirically derived via MEME^35^ from native promoters in a subset of reference genomes per genus, by predicting high-expression candidates with TXpredict^36^ (See Methods). Finally, sequences meeting both criteria were prioritized based on low perplexity scores computed by the Evo 1 8k base model. Language model log-likelihoods were used as robust zero-shot predictors of biological activity^15,25^. This process identified canonical promoter-like motifs in 134 of 288 *Bacteroides* candidates (the −7/−33 sigma factor motif^37^) and in 19 of 453 *Bifidobacterium* candidates (the −10 TAnnnT element^38^) (**Fig. 5b**). For both genera, the top five motif-positive sequences with the lowest perplexity scores were selected for functional characterization. Conversely, since the predicted *Lactobacillus* motif was detected in only two generated sequences (PDiffu_LP_1 and PDiffu_LP_2), we supplemented these two sequences with the three lowest-perplexity candidates (PDiffu_LP_3, PDiffu_LP_4 and PDiffu_LP_5) that passed only the novelty filter.

**Fig 5.**
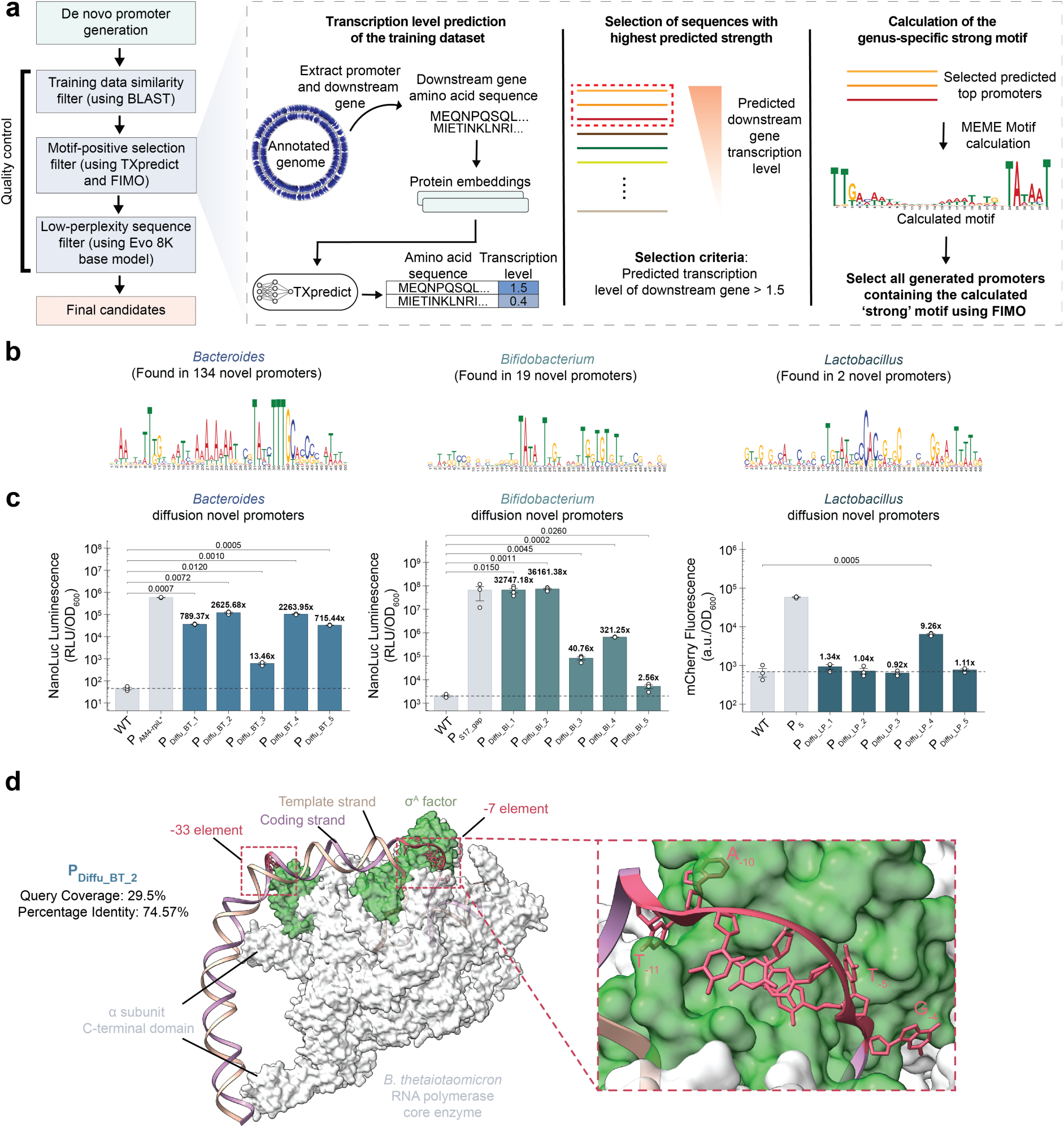
Functional validation of diffusion-generated promoters in non-model gut bacteria selected through our filtering pipeline. **a,** Pipeline for motif-based filtering of generated promoters. **b,** Representative strong motifs identified for each tested non-model bacterial genus. **c,** Reporter activity (fold-change over wild-type, bold) of five computationally prioritized synthetic promoters tested individually in *B. thetaiotaomicron*, *B. infantis* and *L. plantarum*, with native reference promoters (P_AM4-rpiL*_, P_S17_gap_, and P_5_) shown for comparison. Bars show mean ± SE; white circles represent individual biological replicates (n = 3 per group, except for P_Diffu_BI_1_, P_Diffu_BI_2_, P_Diffu_BI_3_, P_Diffu_BI_5_ where n = 4 per group). **d,** AlphaFold3 Structural model of the *B. thetaiotaomicron* RNAP holoenzyme comprising the core enzyme (gray) and σ^A^ factor (green) in complex with the de novo-designed promoter P_Diffu_BT_2_. The −7 and −33 promoter recognition elements are highlighted in red.

For all three hosts, this small experimental set yielded functional promoters (**Fig. 5c**). In *B. thetaiotaomicron*, all five synthetic *Bacteroides* promoters were active, although their activities were weaker than that of the strong native PAM4_rpiL* reference. All five *Bifidobacterium* designs were active in *B. infantis*, with PDiffu_BI_1 and PDiffu_BI_2 approaching the strength of the native promoter PS17_gap. In *L. plantarum*, only PDiffu_LP_4 (selected by low perplexity) showed significant activity, albeit below that of the native P5 control, while motif-selected candidates failed. Thus, after testing only five candidates per non-model host, we conclude that a sequence-only generation model combined with in silico prioritization produced functional *cis*-regulatory elements.

The capacity of the diffusion pipeline to capture complex rules was particularly evident in the strong *Bacteroides* promoters (P_Diffu_BT_1_, P_Diffu_BT_2_, P_Diffu_BT_4_, P_Diffu_BT_5_; **Extended Data Fig. 5a,b**), given their unusual promoter architecture. AlphaFold3^29^ modeling of P_Diffu_BT_2_ bound to the *B. thetaiotaomicron* RNAP-σ^A^ holoenzyme (ipTM = 0.86, pTM = 0.87; **Fig. 5d**) predicted the concurrent σ^A^ engagement at the −33 and −7 elements. This interaction featured coding-strand nucleotides A_-10_ and T_-5_ flipping into complementary pockets, a ∼15-nt transcription bubble initiating upstream of the −7 element, and putative UP-element interactions driven by RNAP α-CTD binding to an upstream AT-rich tract.

We observed a similar pattern for the strong *Bifidobacterium* promoters (P_Diffu_BI_1_ and P_Diffu_BI_2_; **Extended Data Fig. 6a,b**). Structural modeling of P_Diffu_BI_2_ with the *B. infantis* RNAP-σ^70^ holoenzyme (ipTM = 0.82, pTM = 0.84; **Extended Data Fig. 6c**) revealed canonical −35 and −10 recognition, a ∼17-nt transcription bubble, and analogous α-CTD upstream interactions. This sequence distinctness and motif utilization also extended to our top *Lactobacillus* candidate, P_Diffu_LP_4_ (**Extended Data Fig. 7a,b**).

Structural and functional analyses confirm that our generative pipeline, coupled with in silico filtering, produces de novo functional promoters with activity matching that of native sequences across canonical and noncanonical transcription systems. Designed promoters recapitulate key initiation features and system-specific interaction patterns required for productive RNAP engagement.

Many of these synthetic sequences lack detectable homology to natural promoters. To elucidate broader design principles, such as sequence motifs and spatial architectures, we adapted a transcriptional reporter assay for non-model hosts, building on established targeted sequencing methodologies^4^ (**Fig. 6a**; see Methods for details). In this high-throughput assay, a barcoded library places each synthetic promoter upstream of a reporter gene, with a unique barcode in the 3′ untranslated region (UTR). Promoter activity (*T_i_*) is quantified by sequencing barcode abundance via next-generation sequencing (NGS) in the RNA pool and then normalized against its representation in the plasmid DNA library.

**Fig 6.**
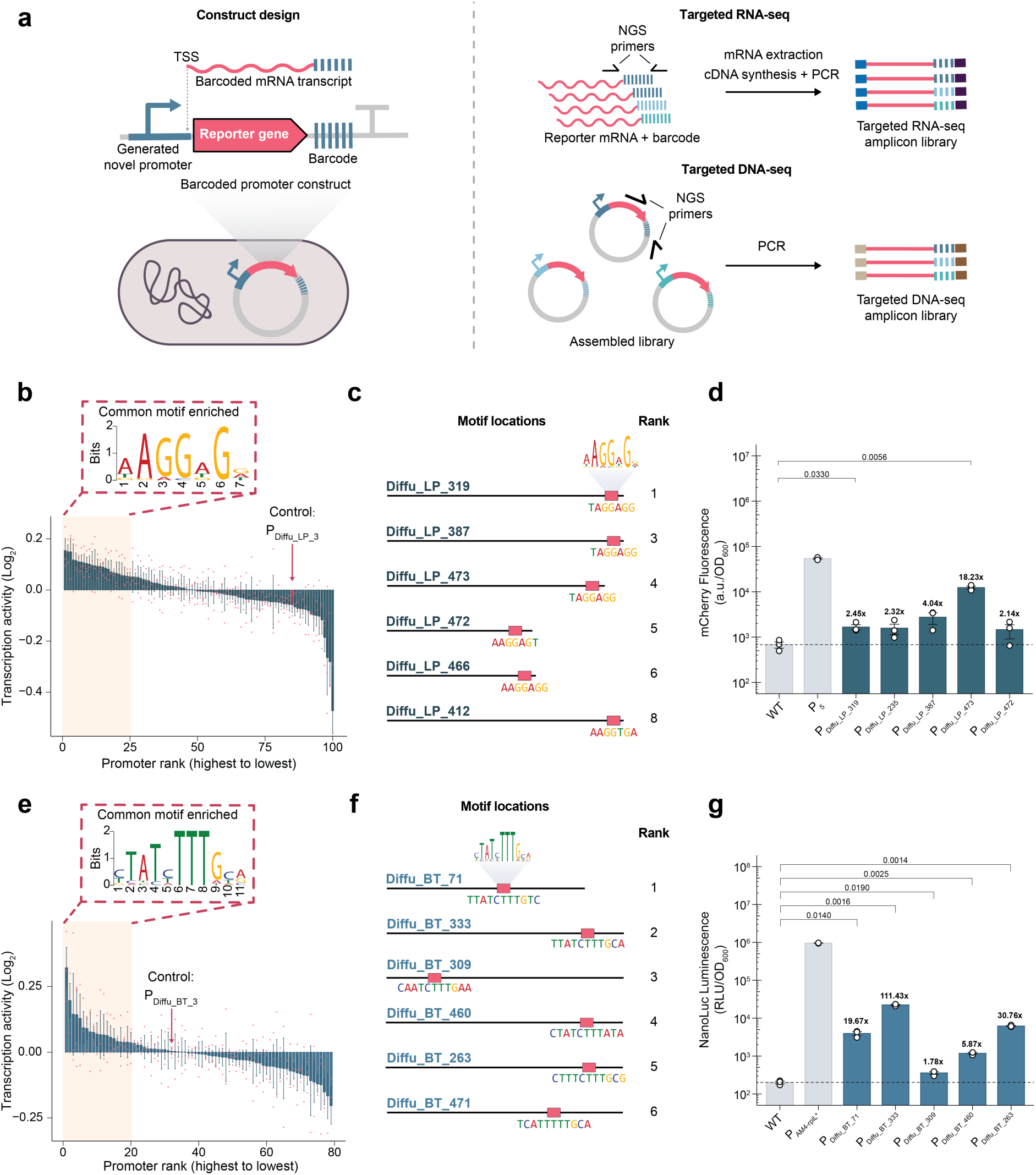
High-throughput characterization of synthetic promoters in non-model gut commensals. **a,** Barcoded reporter strategy for high-throughput and multiplexed promoter activity measurements using targeted RNA and DNA sequencing. **b,** Transcriptional activity of 100 synthetic *Lactobacillus* promoters in *L. plantarum*; the top 25 sequences (shown in yellow) were used for motif discovery; the negative control was P_Diffu_LP_3_ (shown in red). **c,** Motif locations within selected *Lactobacillus* high-activity promoters. **d,** Reporter output of the top five *Lactobacillus* promoters in *L. plantarum*. **e**, Transcriptional activity of 79 synthetic *Bacteroides* promoters in *B. thetaiotaomicron*; the top 20 sequences (shown in yellow) were used for motif discovery; low-expression control P_Diffu_BT_3_ (shown in red). **f,** Motif locations within selected *Bacteroides* high-activity promoters. **g,** Reporter output of the top five *Bacteroides* promoters in *B. thetaiotaomicron.* Bars show mean ± SE; individual dots represent individual biological replicates (n = 3 per group). Statistical significance was determined by an unpaired, two-sided Welch’s t-test.

To validate our high-throughput characterization platform in a non-model host, we screened 100 synthetic *Lactobacillus* promoters in *L. plantarum*. These promoters were prioritized by low Evo 1 8k perplexity. Forty-eight of the promoters screened were functionally active *(log*_2_*T_i_*> 0), and the library displayed a broad dynamic range of transcriptional activities (**Fig. 6b** and **Extended Data Fig. 8a**). The reliability of the assay was confirmed by a baseline negative control (PDiffu_LP_3, *log*_2_*T_i_* ≈ 0) and strong biological replicate correlation (**Extended Data Fig. 8b**). The sequence grammar driving expression was then decoded by de novo motif discovery. A conserved Shine-Dalgarno hexamer (AAGGAG; E-value: 2.8×10⁻⁶) was found for the top 25 variants identified, precisely 5-8 bp upstream of the start codon (**Fig. 6c** and **Extended Data Fig. 8c**), mirroring the usual spatial positioning of promoters and ribosome binding site in *L. plantarum* and other prokaryotes^39,40^. The spontaneous emergence of this motif, as well as its position, indicates that the model had autonomously learned both the necessity of ribosomal recruitment and its requisite spatial syntax.

Individual testing of five high-ranking *Lactobacillus* candidates from this screen confirmed two functionally active promoters, PDiffu_LP_319 and PDiffu_LP_473 (**Fig. 6d**). This discrepancy between the initial high-throughput transcription-based assay and our individual validation aligns with known transcription-translation decoupling^4^, highlighting the influence of post-transcriptional determinants. Heterologous testing in *B. thetaiotaomicron* indicated that cross-host activity was heterogeneous: PDiffu_LP_235, although natively inactive in *L. plantarum*, unexpectedly drove robust heterologous expression in *B. thetaiotaomicron* (**Extended Data Fig. 8d**). This off-target activation, indicating that cryptic elements may opportunistically recruit distantly related transcriptional machinery, illustrates the evolutionary plasticity of bacterial gene expression.

We similarly profiled 79 synthetic *Bacteroides* promoters in *B. thetaiotaomicron* (**Fig. 6e)**. Similar to the *Lactobacillus* library, we observed a broad dynamic range (**Extended Data Fig. 9a**), with biological triplicates showing consistent trends across measurements (**Extended Data Fig. 9b**). Motif analysis of the top 20 active promoters identified the enriched consensus sequence TATnTTTG (E-value: 8.1×10^-3^), which matched previously described canonical *Bacteroides* -7 elements^37^. While the presence of this motif was anticipated, given that our preliminary filter identified it in 134 of the 288 generated candidates (**Fig. 5b**), the top variants exhibited strict sequence conservation and native-like 3′ end localization (**Fig. 6f** and **Extended Data Fig. 9c**).

Unlike the *Lactobacillus* candidates, all five top-ranking *Bacteroides* promoters drove strong activity during native validation, though their activity rankings also mildly deviated from transcriptional predictions. The strongest expression was observed with PDiffu_BT_333 (111.43-fold increase; **Fig. 6g**). When these promoters were transferred to *L. plantarum*, they showed little activity (**Extended Data Fig. 9d**). This robust orthogonality confirms that the model had captured genus-specific regulatory grammar, yielding highly active and taxonomically restricted promoters.

Together, these experiments show that diffusion adaptation of a pretrained genomic foundation model can generate regulatory DNA that is active in phylogenetically distant bacteria. Notably, computational prioritization concentrated initial experimental effort on only five candidates per non-model host, while multiplexed profiling provided a complementary route to interrogate larger design sets. This framework therefore couples generative breadth with selective experimental testing, bypassing the traditional reliance on large-scale synthesis as the primary pathway to functional discovery.

## Discussion

Genomic foundation models learn biological sequence representations from large-scale genomic data. To date, however, these models have primarily been developed as tools for sequence understanding, annotation, and prediction.

Our study demonstrates that pretrained foundation models can also serve as practical engineering platforms. By efficient fine-tuning, we adapted Evo 1 as a diffusion-based gLM and showed that the biological knowledge acquired during large-scale pretraining can be repurposed for programmable *cis*-regulatory DNA engineering without repeating foundation-scale training. More broadly, this work suggests that the next stage of biological foundation models will be defined by their ability to generate functional biological components.

Our results also motivate a shift in generative modeling paradigms for regulatory DNA. While most genomic foundation models adopt AR generative paradigm established in natural language processing, regulatory DNA function depends on combinatorial interactions among sequence elements distributed across an entire regulatory region. This organization motivates generative approaches that can model and refine sequence context bidirectionally rather than generating DNA exclusively through left-to-right factorization. Diffusion-based generation, alternatively, iteratively refines complete sequences in a global context. In our study, sequence characterization indicated that the model learned the spatial grammar needed to recruit species-specific transcriptional machinery, rather than isolated motifs. Although studies of additional regulatory systems and architectures will be required to define the relative strengths of AR and diffusion-based approaches, our findings suggest that biologically motivated generative paradigms may provide new opportunities for regulatory sequence engineering.

Our study also indicates that genetic toolkits can be expanded for non-model microorganisms. Gut commensals are increasingly being engineered as diagnostic biosensors and microbial cell factories, yet few well-characterized regulatory elements are available. For many species, genome sequences are now readily available, whereas experimentally validated promoters remain scarce. Our DAPE-PD framework requires only publicly available genomic sequences for model adaptation and can generate and prioritize promoter candidates without large functional training datasets. Because computational prioritization can substantially reduce the experimental search space, this strategy facilitates the development of regulatory parts in emerging microbial chassis and may accelerate the broader adoption of non-model bacteria in biotechnology.

Our study focused on bacterial constitutive promoters and evaluated representative gut-associated species, but additional regulatory elements and phylogenetically diverse microorganisms remain to be explored. Our results establish a practical framework for converting pretrained genomic foundation models into programmable engineering platforms, bridging advances in artificial intelligence with the development of genetic toolkits for synthetic biology and living medicines. Future work could extend this framework to other classes of *cis*-regulatory sequences, integrate iterative experimental feedback through active learning, and adapt increasingly powerful genomic foundation models as they become available.

## Methods

### Model Architecture

Our model architecture is based on the StripedHyena backbone introduced in Evo 1^41^. The model comprises 32 stacked blocks with a hidden dimension of 4096. Each block consists of a sequence-mixing layer, responsible for modeling dependencies along the sequence dimension, followed by a channel-mixing layer operating along the feature dimension.

Within the sequence-mixing layers, the architecture interleaves 29 Hyena layers with 3 self-attention layers placed at equal intervals. The self-attention layers adopt the standard multi-head attention formulation and use rotary position embeddings (RoPE)^42^ to encode relative positional information. Unlike the original Evo 1 model, which employs a causal attention mask for autoregressive generation, we removed the causal mask during fine-tuning, enabling fully bidirectional attention to support masked diffusion training.

The Hyena layers implement input-dependent sequence mixing through a combination of short convolutions (gating mechanisms) and implicit long convolutions, providing a subquadratic alternative to self-attention for long-range dependency modeling. In Evo 1, causality is enforced by parameterizing strictly causal convolutional filters. In contrast, to enable bidirectional information flow, we modified the Hyena convolution operators when using FFT-based convolution algorithms by mirroring the original left-to-right causal filters, thereby incorporating right-to-left contextual information and ensuring symmetric, bidirectional sequence modeling.

DNA sequences are tokenized at single-nucleotide resolution. The model uses an effective vocabulary of four tokens, corresponding to the four nucleotide bases, drawn from a total vocabulary of 512 UTF-8 characters, which allows flexibility for incorporating additional special tokens during fine-tuning and generation.

### Masked Diffusion Modeling

Masked diffusion models, designed for modeling discrete data^43^ such as text, are directly applicable to DNA sequences, which consist of a small categorical vocabulary of four nucleotides. In diffusion models, a forward process *q* gradually corrupts clean data *x*_0_ into a noisy variable *x_t_*, while a learned reverse process *p_θ_* parameterized by *θ* aims to denoise *x_t_* to reconstruct *x*_0_. In masked diffusion, corruption is implemented by mapping tokens into an absorbing state represented by a special [MASK] token. Given a continuous diffusion time *t* ∈ (0,1), we randomly replaced each token in the input sequence with [MASK] using a mask ratio proportional to *t*. Larger values of *t* therefore corresponded to more heavily corrupted sequences. Training followed the standard diffusion variational formulation. After deriving the evidence lower bound (ELBO) for the masked diffusion process, the objective reduces to a reconstruction loss applied only at masked positions. Concretely, we optimized the following loss:

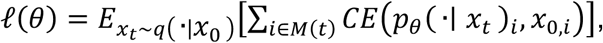

where *M*(*t*) is the set of masked positions and CE denotes cross-entropy.

During generation, we initialized the sequence entirely with [MASK] tokens and performed iterative denoising. At each iteration, the model predicted token identities for all masked positions using *p_θ_*(*x*_0_|*x_t_*), and a randomly selected subset of positions was recovered, while the remaining positions stayed masked and were updated in subsequent iterations. The random selection at each step introduces stochasticity into the generation process, allowing the masked diffusion model to produce diverse samples rather than converging to a single deterministic sequence.

Training a masked diffusion model from scratch typically requires large-scale pretraining. To reduce this cost, we leveraged the pretrained autoregressive Evo 1 model and fine-tuned it into a masked diffusion model. Specifically, we inherited the model parameters from Evo 1, switched the architecture to support bidirectional context, and optimized the model using the masked diffusion objective described above. This adaptation allows the model to retain the sequence representations learned during Evo 1 pretraining while enabling diffusion-based generation.

To support conditional generation across a variety of bacterial species, we prepended the species name, encoded using standard UTF-8 characters, to the promoter sequence during training. At inference time, providing a species prefix allows the model to generate promoter sequences conditioned on the specified species.

### Training Procedure

All models were trained on 8 NVIDIA H100 GPUs using bfloat16 (BF16) precision and ZeRO Stage-2 optimization. The per-GPU batch size was set to 16. Promoter sequences are relatively short; therefore, all input sequences were padded to a fixed length of 220 tokens. We used the character “X” as the [MASK] token, which was not updated during the original autoregressive pretraining of Evo 1 and was only introduced and optimized during diffusion-based fine-tuning. Optimization was performed using the Adam optimizer with a maximum learning rate of 3e-4. Training employed a linear warmup over the first 20,000 steps, followed by a cosine decay learning rate schedule.

For *Bacillus, Pseudomonas,* and *Escherichia* datasets, models were trained for 3 epochs. For non-model bacterial species, training was extended to 100 epochs to compensate for the increased data heterogeneity and the reduced dataset size. The label-shift operation from autoregressive pretraining was retained to preserve alignment with the pretrained hidden representations. During generation, we generated 500 samples with 220 lengths, where the diffusion timesteps equal to generation length.

### Intergenic Region Mining Pipeline

To obtain the fine-tuning dataset of the models, we began by retrieving all available annotated and complete genomes for the genus of interest from the National Center for Biotechnology Information (NCBI) database using the NCBI Datasets command-line interface. Intergenic sequences were subsequently extracted from each genome using custom Python scripts. Specifically, these scripts identified intergenic regions located upstream of the start codon of the downstream gene and retained only those sequences longer than 50 bp. For sequences exceeding 200 bp, we trimmed each to a uniform length of 200 bp, starting immediately upstream of the start codon.

To mitigate redundancy and preferentially retain larger, more complete promoter sequences over short intergenic fragments from the metagenomic mining dataset, we performed sequence clustering using the Linclust algorithm from the MMseqs2 suite^26^. The clustering was implemented with the following criteria: (1) the shorter sequence in any pairwise comparison must align over its entire length to the longer sequence, (2) the alignment must exhibit at least 95% sequence identity, and (3) the longest sequence within a cluster is selected as the representative. These parameters were specified in Linclust as follows: --kmer-per-seq 80 -c 1.0 --cluster-mode 2 --cov-mode 1 --min-seq-id 0.95. The final fine-tuning dataset was constructed by extracting these representative sequences and subsequently filtering out any sequences containing non-nucleotide characters.

### Bacterial Strains, Growth Conditions, and Expression Vectors

All bacterial strains and plasmids used in this study, together with the sequences of the primers and genes, are detailed in **Supplementary Table 1** and **Extended Data Fig. 10**. Routine cloning was performed using *E. coli* TOP10 and 10-Beta, whereas *E. coli* S17-1 λpir was utilized for *Bacteroides* plasmid assembly and plasmid conjugation experiments. *Escherichia* promoter assays were conducted in *E. coli* Nissle 1917. All *E. coli* strains were cultured aerobically in Luria–Bertani (LB) broth at 37°C with shaking (250 rpm); plasmid-bearing strains were selected with kanamycin (50 μg/mL). *L. plantarum* NC8 was cultured aerobically in de Man, Rogosa and Sharpe (MRS) medium at 30°C without shaking; plasmid selection was maintained with erythromycin (10 μg/mL). *B. thetaiotaomicron* VPI-5482 was cultured statically and anaerobically inside a vinyl anaerobic chamber (Coy Laboratory) at 37°C in pre-reduced brain heart infusion–supplemented (BHIS) medium containing hemin (5 µg/mL), vitamin K (1 µg/mL), and cysteine (0.05%), hereafter referred to as BHIS (+Hemin, +VitK, +Cys). Selective media for *B. thetaiotaomicron* plasmid constructs were supplemented with gentamicin (200 μg/mL) and erythromycin (25 μg/mL). *B. infantis* S12 was cultured statically and anaerobically at 37°C in pre-reduced MRSc medium (MRS supplemented with 0.5 M sucrose and 0.05% L-cysteine); plasmid constructs were selected with chloramphenicol (10 μg/mL).

Several reporter plasmids were constructed for each target bacteria. For *Escherichia* promoter constructs, we employed a custom ColE1-*Lpl* rpoD-GFP-mCherry-axe/txe vector. This plasmid carries a GFP reporter gene driven by the PProD constitutive promoter, along with the heterologous sigma factor RpoD from *L. plantarum* to expand the promoter repertoire recognized by the *E. coli* transcriptional machinery^44^ and allow a better assessment of high-throughput library behavior. Additionally, the plasmid includes an mCherry reporter gene downstream of the promoter insertion site and a kanamycin resistance marker. For the *Lactobacillus* promoter constructs, a modified pSIP403^45^ (Addgene plasmid #122028) derivative harboring an mCherry reporter gene and an erythromycin resistance marker was used. For the *Bacteroides* promoter constructs, a modified version of pMM597^5^ (Addgene plasmid #68886) was constructed, containing a NanoLuc reporter gene, an erythromycin selectable marker for *Bacteroides*, and the intN2 tyrosine integrase for site-specific genomic integration. However, given the poor transformation efficiency of this plasmid in *B. thetaiotaomicron*, we adapted a non-integrative pFW2000^46^ derivative (Addgene plasmid #224211), which was modified to carry a NanoLuc cassette and a *Bacteroides* erythromycin resistance gene for use in our transcription-based high-throughput screening. Finally, for the *Bifidobacterium* promoter constructs, we constructed a pKO403^47^ (Addgene plasmid #174725) variant carrying a chloramphenicol resistance marker and a NanoLuc reporter gene.

### *Escherichia* Library Cloning, Transformation and FACS-Seq Experiments

For high-throughput screening, all three generated promoter libraries (Diffusion, Joint AR, and Separate AR) were subjected to an initial bioinformatic filter to remove variants with high sequence homology to known promoters. For this, we defined a "query identity" metric as the number of exact matches of the alignment relative to the length of the query sequence. We retained only those sequences from the generated libraries exhibiting a maximum query identity of < 30% against the training dataset. Subsequently, to enable an unbiased comparison between strategies with unequal library sizes, we used the Promoter Calculator^28^ to predict the relative strength of all filtered variants and selected the top 100 scoring promoters from each library for downstream analysis.

For the pooled FACS screening, two distinct library pools were synthesized. The first pool contained 500 promoters, comprising 188 Joint AR and 132 Separate AR variants that passed the query identity filter (inclusive of the top 100 Promoter Calculator-ranked candidates for both strategies), supplemented with select variants from earlier generation attempts. The second pool consisted exclusively of the top 100 filtered promoters from the Diffusion library. These two pools were synthesized by Integrated DNA Technologies (IDT) using their oligo library synthesis platform. Pooled oligos were PCR-amplified (Phanta UniFi Master Mix, Vazyme) and cloned upstream of the mCherry reporter in the ColE1-Lpl rpoD-GFP-mCherry-axe/txe vector via Gibson Assembly (ClonExpress Ultra One Step Kit V3, Vazyme). Assembled constructs were purified by ethanol precipitation.

To prepare *E. coli* electrocompetent cells, an overnight culture was subcultured 1:100 into 50 mL of fresh LB medium and incubated at 37°C until reaching mid-exponential phase (OD_600_ ∼0.5-0.6). All subsequent harvesting, washing, and aliquoting steps were strictly performed on ice or at 4°C using pre-chilled materials. Cells were chilled on ice for 15 min and then harvested by centrifugation at 4,000 rpm for 15 min at 4°C. To remove residual salts, the pellet was washed sequentially through a series of gentle resuspension and centrifugation steps (4,000 rpm for 15 min at 4°C): first with 50 mL, then 25 mL, 10 mL, and 5 mL of ice-cold sterile water, followed by a final wash in 5 mL of ice-cold 10% (v/v) glycerol. Finally, the cells were resuspended in 500 µL of ice-cold 10% glycerol, divided into 100 µL aliquots in pre-chilled microcentrifuge tubes, and immediately stored at -80°C.

Bacterial cells were transformed with the assembled library constructs by electroporation by adding 100 ng of plasmid library DNA to 100 µL of thawed electrocompetent cells. The mixture was incubated on ice for 25 min and subsequently transferred to a pre-chilled 0.1-cm electroporation cuvette. Electroporation was executed using a Gene Pulser Xcell system (Bio-Rad) using the following parameters: 1.8 kV, 25 µF and 200 Ω. Immediately following the pulse, the cells were resuspended in 900 µL of LB broth and allowed to recover at 37°C with shaking (250 rpm) for 1 h. Following the recovery period, the cultures were plated onto LB-kanamycin agar.

After ∼18 h of incubation at 30°C, library colonies were scraped from the plate provided that the estimated coverage (calculated as the total colony count divided by the library size) exceeded 50-fold. Plasmid DNA was extracted from the pooled harvests using the TIANprep Midi Plasmid Kit (Tiangen). Finally, *E. coli* Nissle 1917 electrocompetent cells were transformed with each plasmid library at a >50-fold coverage following the same electroporation protocol and competent cell preparation procedure described above.

For flow cytometry analysis, glycerol stocks of each *E. coli* Nissle 1917 library were inoculated into 25 mL of LB-kanamycin medium in a 1:100 ratio and incubated overnight at 30°C with shaking at 250 rpm. The following day, the overnight cultures were subcultured 1:100 in fresh LB with kanamycin and grown at 37°C, 250 rpm until they reached mid-exponential phase (OD_600_ ∼0.3-0.4). The cells were then pelleted by centrifugation at 6,000 rpm for 15 min at 4°C, washed with an equal volume of cold PBS, and finally diluted in cold PBS to a final concentration of approximately 6 × 10⁶ cells/mL, based on an estimated conversion of OD_600_ = 1.0, corresponding to 8 × 10^8^ cells/mL.

Flow cytometry analysis and cell sorting were performed on a BD FACSAria III Cell Sorter (BD Biosciences). A custom gate arrangement was established to span the full dynamic range of the mCherry signal (PE-Texas Red-A). The subpopulation of cells exhibiting the highest fluorescence intensity (∼1000 events) was collected via sorting into a tube containing 500 µL of LB medium. The sorted population was subsequently incubated in 10 mL of LB-kanamycin medium at 30°C, 250 rpm for ∼18 hours, after which plasmid DNA was extracted from the culture using a TIANprep Midi Plasmid Kit (Tiangen).

To quantify the enrichment of promoter variants, plasmid DNA extracted from the sorted populations was subjected to NGS amplicon sequencing. Promoter regions were PCR-amplified (15 cycles, NEBNext Ultra II Q5 Master Mix, New England Biolabs) with custom primers containing Illumina-compatible adapters and unique dual indexes (i5 and i7), after which the band corresponding to the size of the amplified promoters was purified from the reaction mix using MagicPure Size Selection DNA Beads (TransGen Biotech). The pooled amplicon libraries were sequenced on a NovaSeq X Plus platform (Illumina) using a PE-150 paired-end configuration. The resulting sequencing reads were demultiplexed and aligned to the reference library of promoter sequences using BWA-MEM^48^ to quantify the counts of each variant.

To maintain a rigorously equalized sample size for functional evaluation, we focused our downstream analysis of the combined AR library on only the top 100 Promoter Calculator-ranked variants from the screened Joint AR and Separate AR pools, matching the scale of the Diffusion library (n=100). Consequently, all comparative analyses are based on the equal subset of 100 filtered promoters per generation strategy. To determine whether a specific promoter was enriched in the harvested high-fluorescence population, we established a strict activity threshold requiring a minimum read count of 1,000 in the sorted bin.

FACS-seq hits were individually validated by first inserting the promoter sequences into the ColE1-Lpl rpoD-GFP-mCherry-axe/txe reporter plasmid upstream of the mCherry gene via Golden Gate Assembly using PaqCI (New England Biolabs) restriction sites introduced by PCR. Electrocompetent *E. coli* Nissle 1917 was then transformed with the assembled plasmids. Transformants were selected on LB-kanamycin agar incubated overnight at 37°C. Single colonies were then used to inoculate 5 mL of LB-kanamycin medium; cultures were grown overnight at 37°C and 250 rpm, diluted 1:100 into fresh selective medium, and grown under the same conditions to mid-exponential phase (OD_600_ ∼0.3–0.4). mCherry fluorescence was then quantified using a BioTek Synergy H1 microplate reader (Agilent).

### Filtering Pipeline for Promoters Generated for Non-model Bacteria

To overcome the inherent challenges of performing high-throughput promoter screening in non-model bacterial species, we tried to develop a systematic pipeline for preselecting a subset of generated promoter sequences most likely to function effectively when experimentally validated in living cells.

In the first filtering stage (novelty-based filtering), we assessed the generated promoter sequences for redundancy by aligning them against their respective fine-tuning datasets using BLAST. Sequences with query coverage of 30% or higher against the fine-tuning dataset were excluded to favour sequence novelty.

The second stage (motif-based filtering) aimed to identify sequence motifs that might be associated with strong promoter activity in the reference genomes used for model training. We hypothesized that predicted downstream gene transcription levels could serve as a proxy for promoter strength. For each non-model bacterial dataset, we considered a subset of genomes for analysis: 536 *Lactobacillus*, 496 *Bifidobacterium*, and 193 *Bacteroides* genomes. For each mined promoter sequence from the non-model bacterial genomes, we also extracted the downstream gene and estimated its transcriptional activity using TXpredict^36^, which outputs a z-score–normalized predicted expression level. Intergenic regions with a downstream predictive score greater than 1.5 (i.e., more than 1.5 standard deviations above the mean) were designated as strong promoter candidates. These regions were then analyzed using MEME^35^ to derive 50 bp consensus motifs, and we subsequently retained all generated promoters from the first filter that contained these motifs, identified using FIMO^27^.

In the final stage (gLM-based filtering), we evaluated the plausibility of the remaining promoter candidates using gLM-based metrics. For *Bacteroides* and *Bifidobacterium*, we calculated the perplexity of each filtered sequence using the Evo 1 8k base model and selected the five promoters with the lowest perplexity values for experimental testing. For the *Lactobacillus* dataset, only two promoters met the criteria of the first two filters; therefore, to ensure a consistent experimental sample size, we additionally selected the top three promoters with the lowest calculated perplexity that passed only the novelty-based filter but not the motif-based one. This yielded five promoters per genus for experimental validation.

### *B. thetaiotaomicron* Conjugation and Assessment of Construct Strength of Selected Promoters

Selected promoter candidates were individually cloned upstream of the NanoLuc reporter gene in plasmid pMM597 via Golden Gate Assembly, using PaqCI restriction sites introduced by PCR. The resulting constructs were transformed into electrocompetent *E. coli* S17-1 λpir (Prepared and transformed as previously mentioned). Transformation of *B. thetaiotaomicron* was performed using a conjugation protocol adapted from a previously described method^5^. Briefly, the pMM597 plasmid carries an intN2 tyrosine integrase that mediates site-specific recombination between the attBT site in the *B. thetaiotaomicron* chromosome and the attN site on the plasmid, enabling stable single-copy integration of the reporter cassette.

For conjugation, donor *E. coli* S17-1 λpir harboring the assembled constructs were grown for 18 h at 37 °C with shaking (250 rpm) in LB-erythromycin (250 μg/mL) broth. Recipient *B. thetaiotaomicron* wild-type cultures were grown anaerobically for 24 h in BHIS (+Hemin, +VitK, +Cys) broth. To initiate mating, 250 µL of donor *E. coli* culture was pelleted by centrifugation at 5,000 × g for 5 min and the supernatant discarded. The pellet was resuspended in 1 mL of *B. thetaiotaomicron* recipient culture, followed by a second centrifugation under the same conditions. The resulting pellet was resuspended in 25 µL of pre-reduced BHIS, spotted onto non-selective BHIS agar plates pre-warmed to 37°C, and plates were incubated overnight under aerobic conditions.

Following conjugation, the spots were scraped with sterile loops, resuspended in 1 mL of BHIS (+Hemin, +VitK, +Cys), and mixed thoroughly by pipetting. Aliquots (250 µL) were plated onto selective BHIS (+Hemin, +VitK, +Cys) agar containing gentamicin and erythromycin. Plates were incubated anaerobically at 37°C for 48 h.

Single colonies from selective plates were restreaked on the same medium to verify purity and remove residual *E. coli*. *B. thetaiotaomicron* transconjugants were cultured in 3 mL BHIS (+Hemin, +VitK, +Cys) broth with gentamicin and erythromycin under anaerobic conditions at 37°C overnight. Glycerol stocks were then prepared for long-term storage.

For luminescence measurements of individual promoter constructs, 2 mL cultures of *B. thetaiotaomicron* harboring the relevant plasmids were collected during mid-exponential phase (OD_600_ ∼0.3–0.4). Cell pellets were obtained by centrifugation at 4,000 rpm for 5 min and resuspended in 200 µL of prewarmed GUS lysis buffer (50 mM sodium phosphate, 5 mM dithiothreitol, and 0.1% Triton X-100) supplemented with lysozyme (20 mg/mL), followed by incubation at 37 °C for 30 min to achieve cell lysis. Luciferase activity was then quantified using the Nano-Glo Luciferase Assay System (Promega) according to the manufacturer’s instructions. Namely, 50 µL of clarified lysate was mixed with an equal volume of Nano-Glo assay reagent, incubated for 5 min at room temperature, and luminescence was measured using a BioTek Synergy H1 microplate reader.

### *B. infantis* Transformation and Assessment of Construct Strength of Selected Promoters

Candidate promoter sequences were individually cloned upstream of the NanoLuc reporter gene in plasmid pKO403 via Golden Gate Assembly, using BsmBI (New England Biolabs) restriction sites introduced by PCR. Electrocompetent cells of *B. infantis* were prepared following a previously described protocol with minor modifications^49^. Briefly, a single colony from a freshly streaked MRSc (MRS + 0.05% L-cysteine hydrochloride) plate was inoculated into MRSc broth and incubated statically at 37°C under anaerobic conditions overnight. The following day, the overnight culture was diluted 1:50 into fresh MRScs broth and grown until the OD_600_ reached 0.15–0.3. Cells were then harvested and washed twice with pre-cooled sucrose-citrate buffer (0.5 M sucrose, 1 mM citrate buffer, pH 6.0) by centrifugation at 3,000 × g for 10 min at 4°C. Finally, the cell pellet was resuspended in sucrose-citrate-glycerol buffer (0.5 M sucrose, 1 mM citrate buffer, 15% glycerol, pH 6.0), aliquoted into 100-μL portions, and stored at -80°C until use.

For electroporation, a frozen aliquot of competent cells was thawed on ice for 10 min and mixed with 500 ng of plasmid DNA, followed by incubation on ice for 30 min. The mixture was transferred to a pre-chilled 0.1-cm electroporation cuvette. Cells were electroporated using a Gene Pulser Xcell system using the following parameters: 2.0 kV, 25 µF and 200 Ω. Immediately after the pulse, cells were recovered in 1 mL of pre-warmed MRSc broth and incubated anaerobically at 37°C for 4–6 h. Recovered cells were then plated onto MRSc-chloramphenicol agar plates and incubated anaerobically at 37°C for 2–3 days until transformant colonies became visible.

For each construct, three to four individual transformant colonies were picked and cultured overnight in MRSc-chloramphenicol broth. The following day, each culture was diluted 1:50 into fresh selective MRSc broth and grown overnight again. From these secondary cultures, 100 μL or 200 μL aliquots were collected at 4 h post-inoculation, corresponding to exponential growth phase (OD_600_ ∼0.5–0.8). The aliquots were loaded into a 96-well plate and pre-cooled on ice inside an anaerobic chamber for 20 min. The plate was then removed, and cells were pelleted by centrifugation at 2,900 rpm for 10 min at 4°C. The supernatant was discarded, and the pellets were resuspended in 100 μL of PBS. The OD_600_ of each resuspension was measured using a plate reader. Cells were then centrifuged again, and the pellets were resuspended in 100 μL of lysis buffer and incubated at 37°C with shaking at 60 rpm for 1 h. Following lysis, the samples were briefly centrifuged at 2,900 rpm for 5 min at RT. A 50-μL aliquot of the clarified supernatant was mixed with freshly prepared Nano-Glo assay reagent and incubated for 5 min at RT, and luminescence was then measured with a Varioskan LUX multimode microplate reader (Thermo Fisher Scientific). Raw luminescence values were normalized to the corresponding OD_600_ reading.

### *L. plantarum* Transformation and Assessment of Construct Strength of Selected Promoters

Candidate promoter sequences were individually cloned upstream of the mCherry reporter gene in plasmid pSIP403 via Golden Gate Assembly, using PaqCI restriction sites introduced by PCR. The resulting constructs were subsequently introduced into *L. plantarum* NC8 by electroporation.

To prepare electrocompetent cells, a single colony of *L. plantarum* NC8 was picked from a freshly streaked MRS agar plate, incubated for 24 h at 30°C, and inoculated into 3 mL of MRS broth. The culture was grown overnight at 30°C without shaking. The following day, 100 µL of the overnight culture was transferred into 5 mL of SGMRS medium (MRS supplemented with 0.75 M sorbitol and 1% glycine) and the tubes were incubated at 30°C until mid-exponential phase (OD_600_ ∼0.3–0.5). All subsequent steps were performed on ice using pre-cooled equipment. The cells were harvested by centrifugation (6,000 rpm, 10 min, 4°C), and the supernatant was discarded. The pellet was washed twice with 2.5 mL of ice-cold SM buffer (952 mM sucrose, 3.5 mM MgCl₂) and finally resuspended in 80 µL of ice-cold SM buffer.

For electroporation, approximately 500 ng of plasmid DNA was mixed with the competent cells, which were incubated on ice for 10 min. The mixture was then transferred to a pre-chilled 0.2-cm electroporation cuvette. Cells were electroporated by using a Gene Pulser Xcell system with the following parameters: 2.0 kV, 25 µF and 400 Ω. Immediately after the pulse, 0.9 mL of MRS broth was added to the cuvette, and the cells were allowed to recover at 37°C for 3 h without shaking. Following recovery, the culture was centrifuged (4,000 rpm, 3 min), the supernatant was reduced to approximately 150 µL, and the pellet was gently resuspended in the remaining medium. The entire volume was spread onto MRS-erythromycin plates and plates were incubated at 30°C for 48 h. Single colonies were then picked and cultured overnight in MRS-erythromycin broth at 30°C without shaking. Glycerol stocks were prepared from these cultures for long-term storage.

For each promoter construct, 4 mL cultures of *L. plantarum* NC8 transformants were grown in selective MRS broth to mid-exponential phase (OD_600_ ∼0.3–0.4). Cells were harvested by centrifugation (4,000 rpm, 5 min) and resuspended in an equal volume of PBS. The resuspended cells were transferred to a black, clear-bottom 96-well plate, and mCherry fluorescence was measured using a BioTek Synergy H1 microplate reader.

### Assembly and Multiplex Characterization of the *Lactobacillus* and *Bacteroides* Non-Model Bacteria Promoter Libraries by DNA-seq and RNA-seq

From the promoter sequences generated for *Lactobacillus* and *Bacteroides*, 100 candidates for *Lactobacillus* and 79 candidates for *Bacteroides* were selected for high-throughput characterization by using the previously described promoter selection pipeline. Each promoter was synthesized upstream of a custom 8-bp barcode, with two PaqCI recognition sites positioned, in opposite orientations, between the promoter and the barcode. The entire construct was flanked by oppositely oriented BsmBI sites (for *Bacteroides* constructs) or BsaI sites (for *Lactobacillus* constructs).

Libraries were assembled in two stages. In the first stage, PCR-amplified promoter fragments were digested and ligated into their respective vectors— pFW2000 for *Bacteroides* using BsmBI, and pSIP403 for *Lactobacillus* using BsaI—using Golden Gate Assembly (New England Biolabs) for 60 cycles, following the manufacturer’s instructions. The assembled plasmid libraries were used to transform chemically competent *E. coli* TOP10 (for the *Lactobacillus* library) or *E. coli* S17-1 λpir (for the *Bacteroides* library), and cells were plated on LB agar supplemented with 10 µg/mL erythromycin (for the *Lactobacillus* library) or 100 µg/mL carbenicillin (for the *Bacteroides* library). Plates were incubated for approximately 18 h at 37°C. Colonies were scraped, and plasmid DNA was extracted with the TIANprep Midi Plasmid Kit (Tiangen) if the transformation yield exceeded a 50-fold threshold.

In the second stage, the extracted plasmid library was digested with PaqCI (New England Biolabs) to introduce the reporter genes—mCherry for the *Lactobacillus* library and NanoLuc luciferase for the *Bacteroides* library—using another 60-cycle Golden Gate Assembly reaction. The resulting constructs were used to transform *E. coli* TOP10 (for the *Lactobacillus* library) or *E. coli* S17-1 λpir (for the *Bacteroides* library), and plasmid DNA was extracted as described above.

The assembled promoter–reporter libraries were subsequently introduced into electrocompetent *L. plantarum* NC8 (as previously described) or into *B. thetaiotaomicron* via conjugation. Transformants were plated on selective media: MRS agar containing erythromycin (10 µg/mL) for the *Lactobacillus* library, or BHIS (+Hemin, +VitK, +Cys) with gentamicin (200 µg/mL) and erythromycin (25 µg/mL) for the *Bacteroides* library. Plates were incubated at 30°C for the *Lactobacillus* library or at 37°C under anaerobic conditions for the *Bacteroides* library for 48 h. Colonies were collected if transformation efficiency exceeded a 50-fold threshold, as defined previously.

To assess promoter strength in the chosen non-model bacteria in a high-throughput manner, we adapted a previously described protocol^4^ to characterize promoter strength, taking a multiplex approach involving targeted RNA-seq and DNA-seq of the constructs.

For targeted RNA-seq and DNA-seq of each of the libraries, overnight cultures were first made from frozen stocks by diluting 1:100 of the thawed frozen stock in 4 mL of MRS + erythromycin (10 µg/mL) for the *Lactobacillus* library or pre-reduced BHIS (+Hemin, +VitK, +Cys), plus gentamicin (200 µg/mL) and erythromycin (25 µg/mL) for the *Bacteroides* library. Cells were cultured overnight as explained previously.

The following day, overnight cultures were diluted 1:100 into 6 mL of fresh growth medium and grown to mid-exponential phase (OD_600_ ∼0.3–0.4).

For RNA extraction, the 2 mL culture aliquot was centrifuged at 12,000 rpm for 2 min at 4°C. The resulting cell pellet was resuspended in 100 µL of TE buffer (10 mM Tris, 0.1 mM EDTA) containing lysozyme (20 mg/mL) and incubated for 10 min to promote cell lysis. Total RNA was then extracted by using the RNAprep Pure Cell/Bacteria Kit (Tiangen) according to the manufacturer’s protocol. The purified RNA was converted to complementary DNA (cDNA) using the iScript cDNA Synthesis Kit (Bio-Rad).

For DNA extraction, the remaining portion of the previously described culture was centrifuged at 7,500 rpm for 10 min at 4°C and the pellet was resuspended in 180 µL of TET buffer (20 mM Tris-HCl, 2 mM Sodium EDTA, 1.2% Triton X-100) containing 20 mg/mL lysozyme and incubated at 37°C for 30 min. Total DNA was then purified from the resulting lysate using the Blood/Cell/Tissue Genomic DNA Extraction Kit (Tiangen) according to the manufacturer’s instructions.

To quantify each promoter variant at both the DNA and RNA levels, NGS amplicon sequencing was performed on the cDNA and the extracted total DNA. Barcode regions were PCR-amplified for 25 cycles using NEBNext Ultra II Q5 Master Mix (New England Biolabs) with custom primers containing Illumina-compatible adapters and unique dual indexes (i5 and i7). Pooled amplicon libraries were sequenced on a NovaSeq X Plus platform (Illumina) with a PE-150 paired-end configuration. Sequencing reads were demultiplexed and low-quality reads were removed with fastp^50^. Barcode sequences were extracted using custom Python scripts.

The transcription level (*T_i_*) of a given promoter construct *i* was calculated according to the following formula:

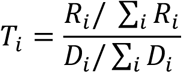

where *R_i_* and *D_i_* represent the read counts corresponding to the RNA and DNA levels, respectively, of each construct.

### Phylogenetic Tree Reconstruction

Phylogenetic relationships were established by tailoring distinct computational pipelines to the genomic architecture of the target taxa. For the model bacterial genera (*Escherichia*, *Pseudomonas*, and *Bacillus*), high-resolution phylogenies were reconstructed using PhyloPhlAn^51^ with the --diversity medium and --fast parameters, leveraging its database of universally conserved prokaryotic marker proteins. For the non-model bacterial genera, we employed a high-fidelity core-genome approach. The pangenome analysis pipeline Panaroo^52^ was utilized to extract core-gene alignments from the curated genomes, and the maximum likelihood phylogenetic trees were subsequently inferred from these core alignments using IQ-TREE^53^. NCBI-reported CheckM completeness scores and species-level taxonomic annotations were projected alongside species-level taxonomic annotations onto the resulting phylogenies.

Finally, viral phylogenies for the genus-specific phages were computed using ViPTreeGen^54^, which constructs robust viral taxonomic trees based on genome-wide sequence similarities. To ensure genomic integrity, phage genome completeness was assessed using the average amino acid identity (AAI) method implemented in CheckV^55^.

## Acknowledgments

We thank Karen Pepper for editing the manuscript. This work was supported by the Research Grants Council of Hong Kong SAR ECS Grant (#26100423 to Y.L.), the Hong Kong Ph.D. Fellowship Scheme (PF22-69790 to M.E.G.), and the Hong Kong University of Science and Technology’s startup grant (R9829 to Y.L.).

