## Supplementary Figures 1-10 for "DAPE-PD: Diffusion adaptation of genomic foundation model enables tailored-generation of *cis*-regulatory DNA"

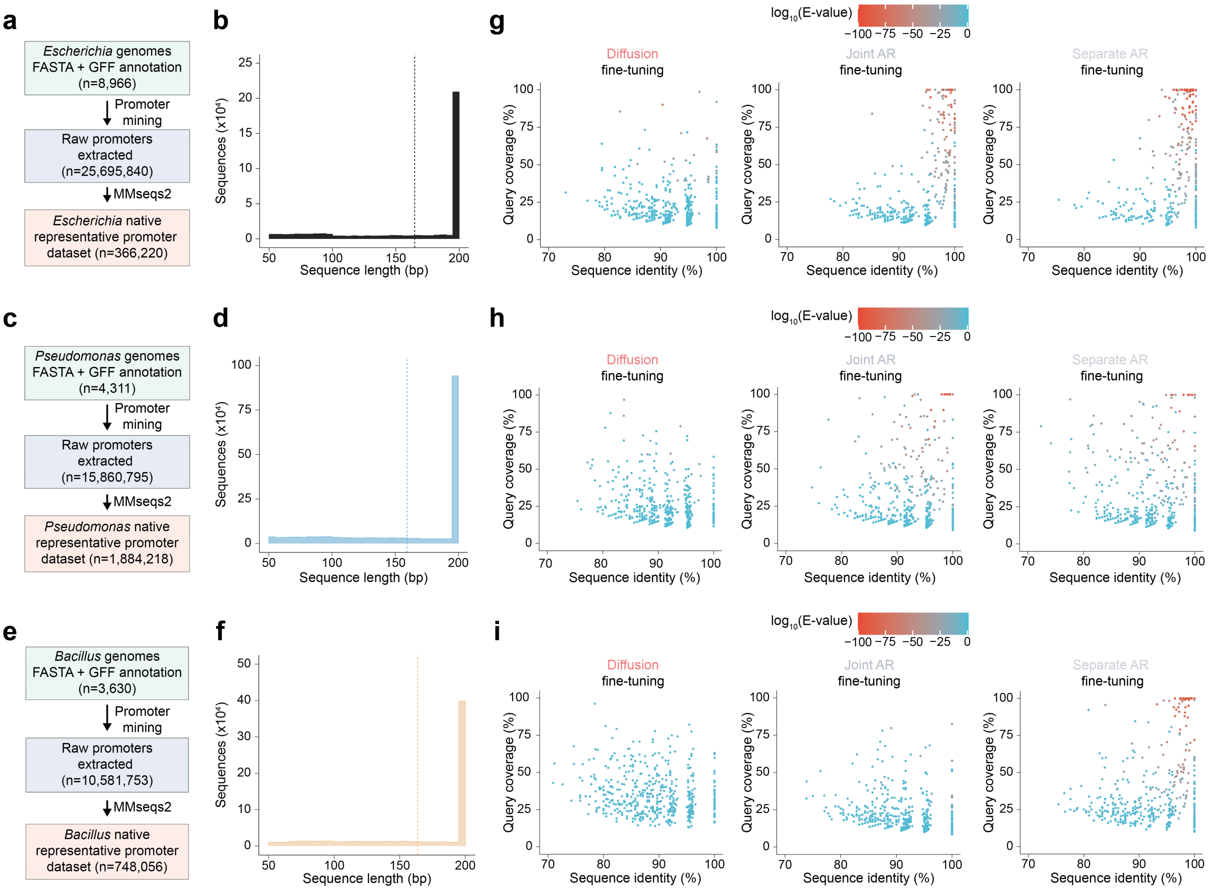


**Extended Data Fig. 1. Mining and generative design of synthetic promoters in model bacteria. a** and **b**, *Escherichia* promoter mining. Workflow for genus-level identification and curation of *Escherichia* promoter sequences (**a**). Length distribution of the curated native *Escherichia* promoter dataset (dotted line, mean) (**b**). **c** and **d**, *Pseudomonas* promoter mining. Workflow for genus-level identification and curation of *Pseudomonas* promoter sequences (**c**). Length distribution of the curated native *Pseudomonas* promoter dataset (dotted line, mean) (**d**). **e** and **f**, *Bacillus* promoter mining. Curation workflow for *Bacillus* promoter sequences (**e**). Length distribution of the curated native *Bacillus* promoter datasets (dotted line, mean) (**f**). **g-i**, Sequence homology of generated sequences for *Escherichia* (**g**), *Pseudomonas* (**h**), and *Bacillus* (**i**).

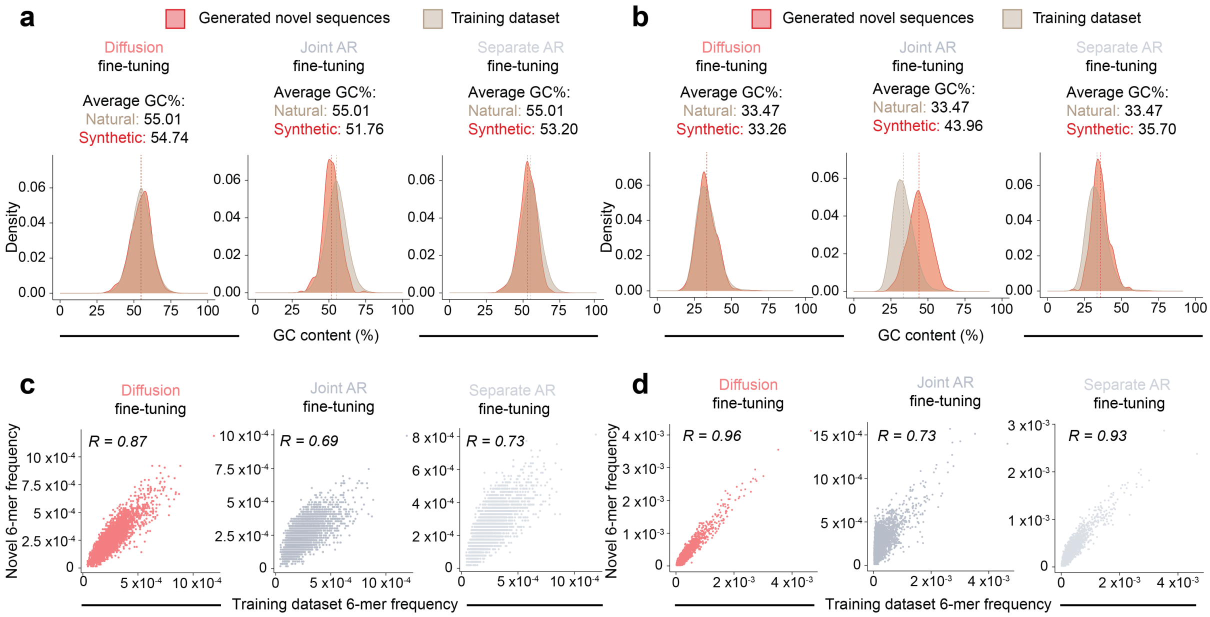


**Extended Data Fig. 2. Characterization of generated novel *Pseudomonas* and *Bacillus* promoters. a-d**, Generated novel promoters exhibit GC-content distributions comparable to those of native sequences (dotted lines indicate sample means) for *Pseudomonas* (**a**) and *Bacillus* (**b**), and maintain strong 6-mer frequency correlations with native profiles for *Pseudomonas* (**c**) and *Bacillus* (**d**). The effect size is reported as Pearson's correlation coefficient.

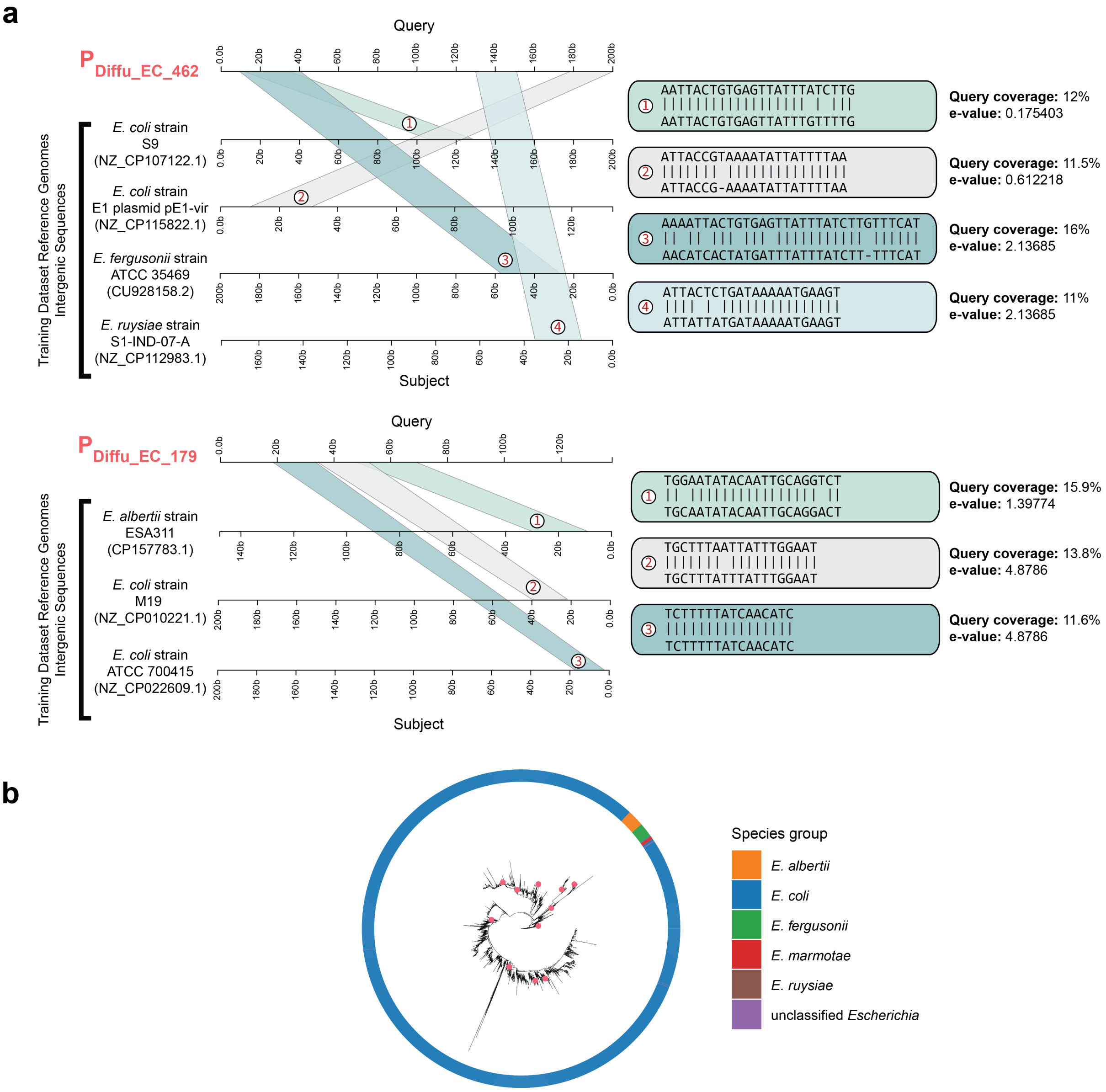


**Extended Data Fig. 3. Functional characterization and sequence analysis of top-performing *Escherichia* promoters identified via FACS-seq. a,** Representative sequence alignments of the top-performing promoters against the training corpus, highlighting the integration of short regulatory motifs derived from diverse genomes. **b,** Phylogenetic distribution of *Escherichia* genomes corresponding to the representative alignments for $P_{Diffu\_EC\_157}$, $P_{Diffu\_EC\_462}$ and $P_{Diffu\_EC\_179}$, highlighted in pink in the phylogenetic tree.

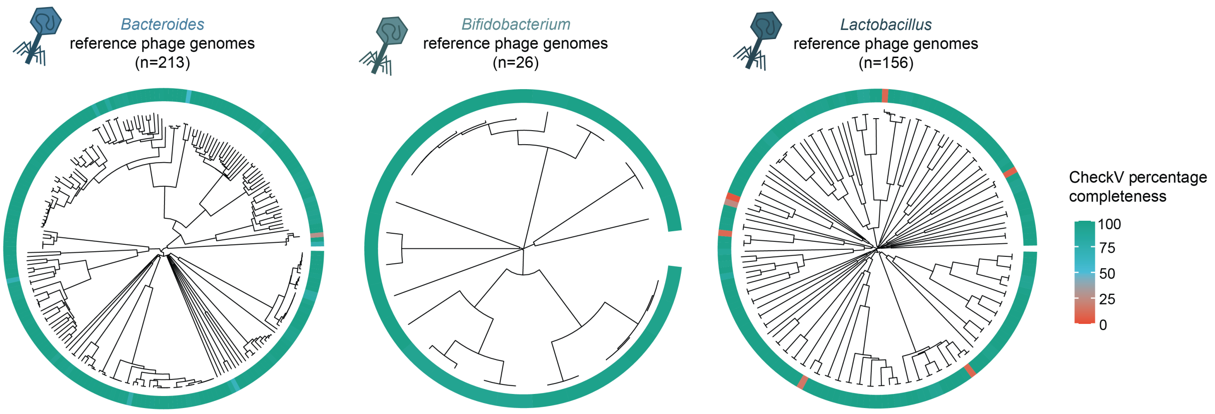


**Extended Data Fig. 4. Phylogeny of phage genomes associated with target non-model bacterial genera.** Phylogenetic trees representing the phage genomes analyzed, corresponding to each non-model bacterial genus of interest. Genome completeness was assessed using the average amino acid identity (AAI) method implemented in CheckV[70].

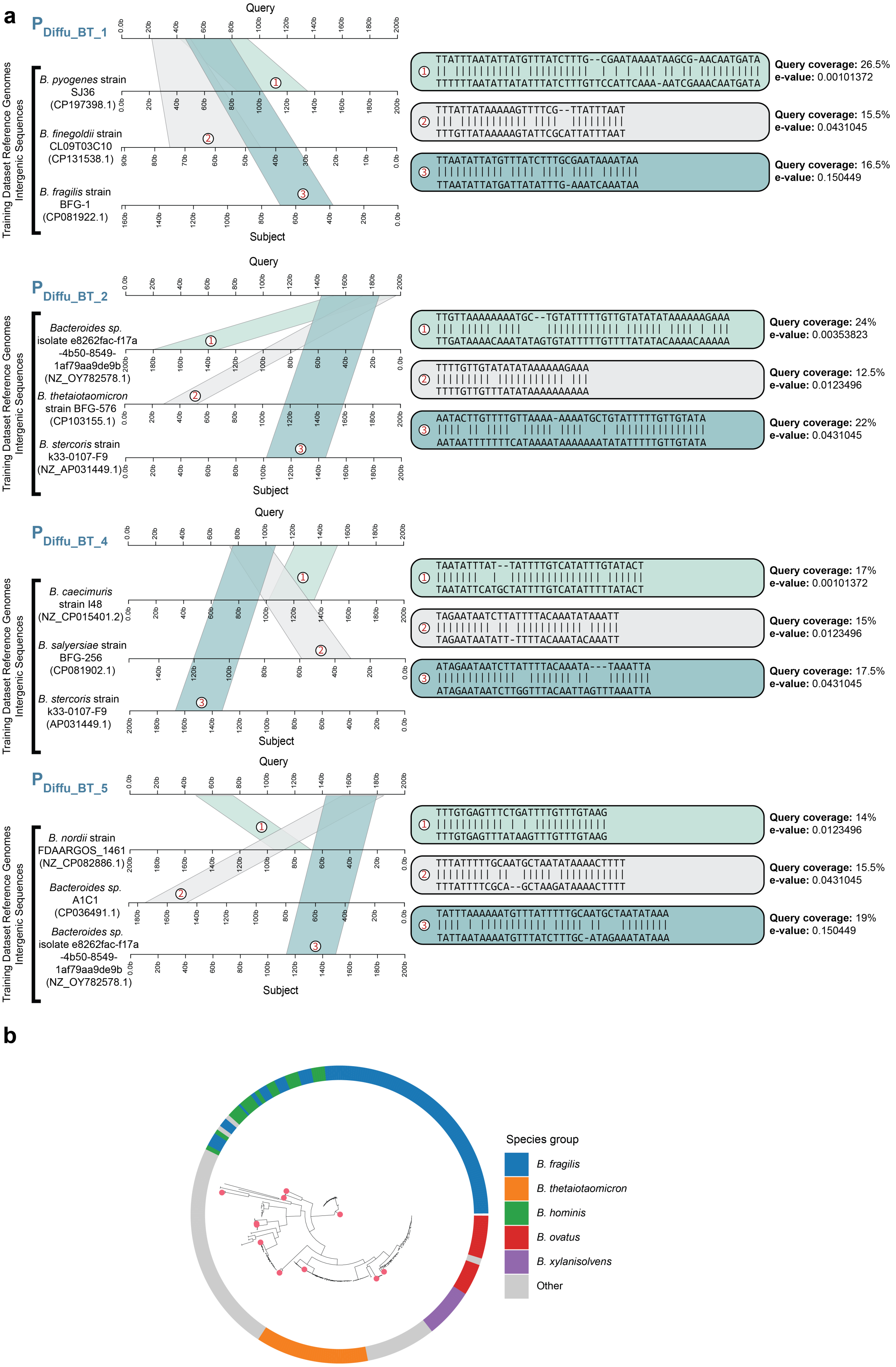

a
P Diffu_BT_1
Query
Subject
Training Dataset Reference Genomes Intergenic Sequences
B. pyogenes strain SJ36 (CP197398.1)
B. finegoldii strain CL09T03C10 (CP131538.1)
B. fragilis strain BFG-1 (CP081922.1)
TTATTTAATATTATGTTTATCTTTG--CGAATAAAATAAGCG-AACAATGATA
TTTTTTAATATTATATTTATCTTTGTTCCATTCAAA-AATCGAAACAATGATA
Query coverage: 26.5%
e-value: 0.00101372
TTTATTATAAAAAGTTTTCG--TTATTTAAT
TTTGTTATAAAAAGTATTCGCATTATTTAAT
Query coverage: 15.5%
e-value: 0.0431045
TTAATATTATGTTTATCTTTGCGAATAAAATAA
TTAATATTATGATTATATTTG-AAATCAAATAA
Query coverage: 16.5%
e-value: 0.150449
P Diffu_BT_2
Bacteroides sp. isolate e8262fac-f17a-4b50-8549-1af79aa9de9b (NZ_OY782578.1)
B. thetaiotaomicron strain BFG-576 (CP103155.1)
B. stercoris strain k33-0107-F9 (NZ_AP031449.1)
TTGTTAAAAAAAATGC--TGTATTTTTGTTGTATATATAAAAAAGAAA
TTGATAAAACAAATATAGTGTATTTTTGTTTTATATACAAAACAAAAA
Query coverage: 24%
e-value: 0.00353823
TTTTGTTGTATATATAAAAAAGAAA
TTTTGTTGTTTATATAAAAAAAAAA
Query coverage: 12.5%
e-value: 0.0123496
AATACTTGTTTTGTTAAAA-AAAATGCTGTATTTTTGTTGTATA
AATAATTTTTTTCATAAAATAAAAAAATATATTTTTGTTGTATA
Query coverage: 22%
e-value: 0.0431045
P Diffu_BT_4
B. caecimuris strain I48 (NZ_CP015401.2)
B. salyersiae strain BFG-256 (CP081902.1)
B. stercoris strain k33-0107-F9 (AP031449.1)
TAATATTTAT--TATTTTGTCATATTTGTATACT
TAATATTCATGCTATTTTGTCATATTTTTATACT
Query coverage: 17%
e-value: 0.00101372
TAGAATAATCTTATTTTTACAAATATAAATT
TAGAATAATATT-TTTTACAAATACAAATT
Query coverage: 15%
e-value: 0.0123496
ATAGAATAATCTTATTTTACAAATA---TAAATTA
ATAGAATAATCTTGGTTTACAATTAGTTTAAATTA
Query coverage: 17.5%
e-value: 0.0431045
P Diffu_BT_5
B. nordii strain FDAARGOS_1461 (NZ_CP082886.1)
Bacteroides sp. A1C1 (CP036491.1)
Bacteroides sp. isolate e8262fac-f17a-4b50-8549-1af79aa9de9b (NZ_OY782578.1)
TTTGTGAGTTTCTGATTTTGTTGTAAG
TTTGTGAGTTTATAAGTTTGTTTGTAAG
Query coverage: 14%
e-value: 0.0123496
TTTATTTTTGCAATGCTAATATAAAACTTTT
TTTATTTTCGCA--GCTAAGATAAAACTTTT
Query coverage: 15.5%
e-value: 0.0431045
TATTTAAAAAATGTTTATTTTTGCAATGCTAATATAAA
TATTAATAAAATGTTTATCTTTGC-ATAGAAATATAAA
Query coverage: 19%
e-value: 0.150449
b
Species group
B. fragilis
B. thetaiotaomicron
B. hominis
B. ovatus
B. xylanisolvens
Other

**Extended Data Fig. 5. Sequence and phylogenetic characterization of *Bacteroides* diffusion functional promoters. a,** Representative sequence alignments of high-performing functional promoters relative to the training corpus. **b,** Phylogenetic distribution of *Bacteroides* genomes corresponding to the representative alignments, highlighted in pink in the phylogenetic tree.

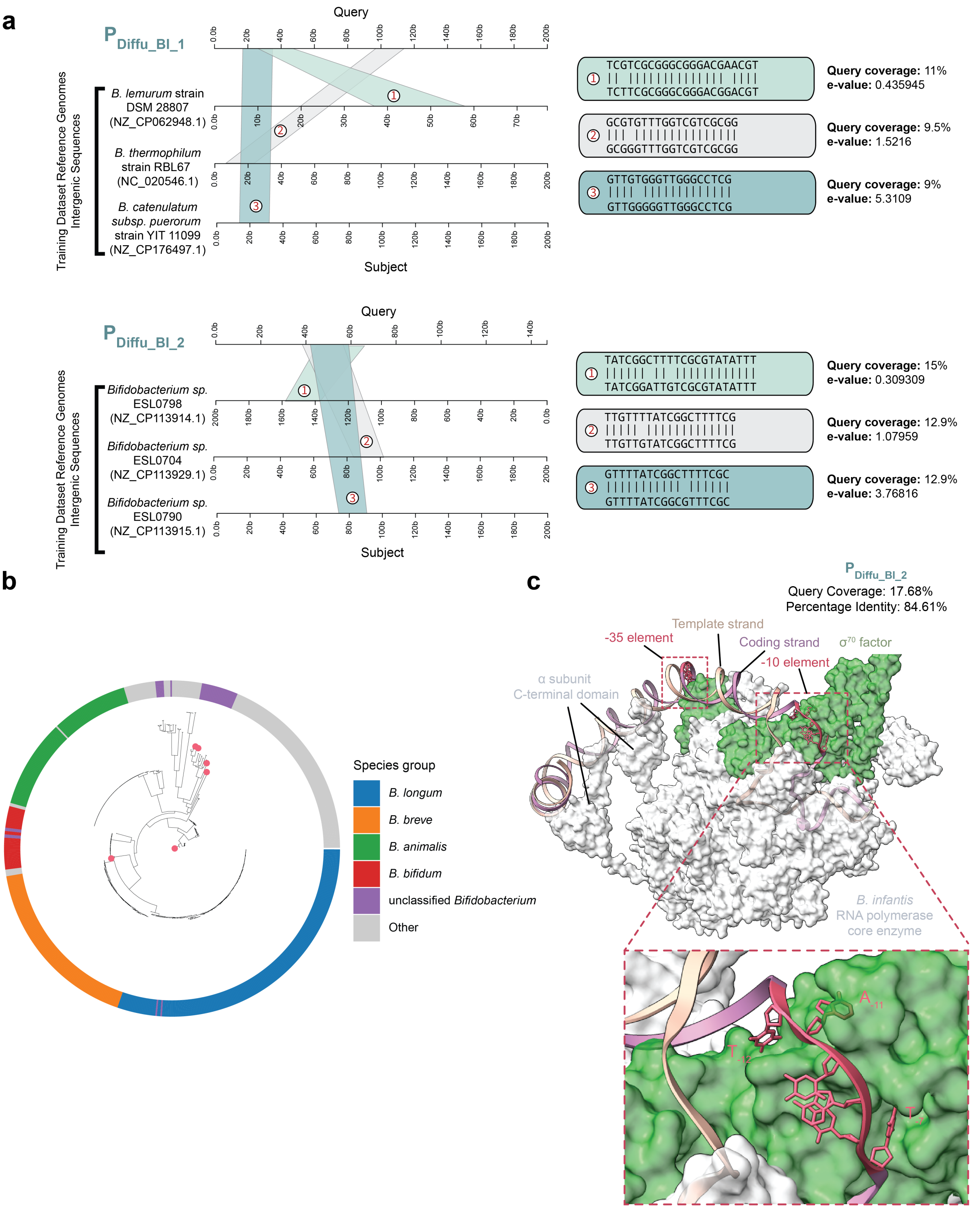


**Extended Data Fig. 6. Sequence and phylogenetic characterization of *Bifidobacterium* diffusion functional promoters. a,** Representative sequence alignments of high-performing functional promoters relative to the training corpus. **b,** Phylogenetic distribution of *Bifidobacterium* genomes corresponding to the representative alignments, highlighted in pink in the phylogenetic tree. **c,** AlphaFold3 Structural model of the *B. infantis* RNAP holoenzyme comprising the core enzyme (gray) and $\sigma^{70}$ factor (green) in complex with the de novo-designed promoter $P_{Diffu\_Bl\_2}$. The −10 and −35 promoter recognition elements are highlighted in red.

**a**

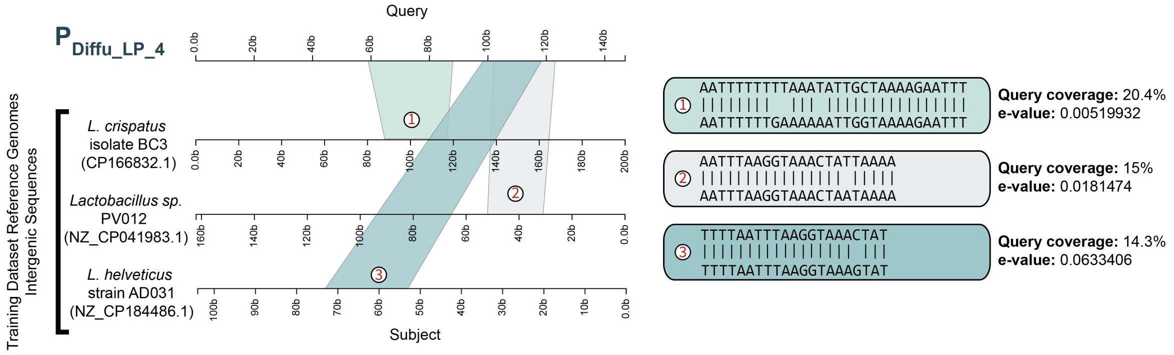


**b**

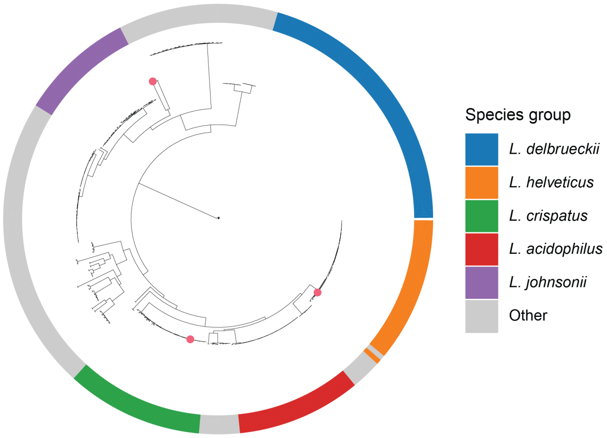


**Extended Data Fig. 7. Sequence and phylogenetic characterization of *Lactobacillus* diffusion functional promoters. a,** Representative sequence alignments of high-performing functional promoters relative to the training corpus. **b,** Phylogenetic distribution of *Lactobacillus* genomes corresponding to the representative alignments, highlighted in pink in the phylogenetic tree.

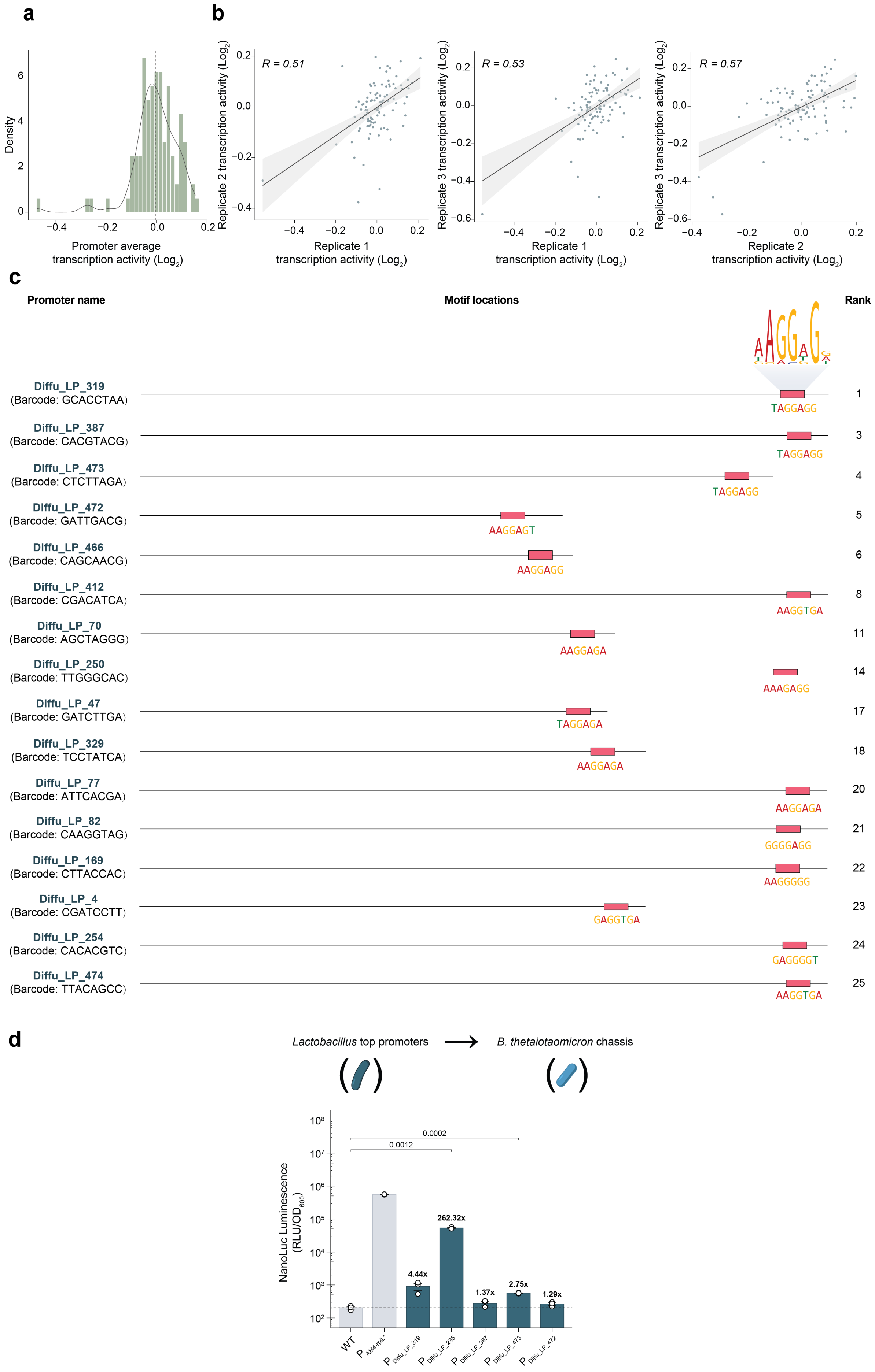

a
Density
Promoter average
transcription activity (Log2)
b
R = 0.51
R = 0.53
R = 0.57
Replicate 2 transcription activity (Log2)
Replicate 3 transcription activity (Log2)
Replicate 1
transcription activity (Log2)
Replicate 2
transcription activity (Log2)
c
Promoter name
Motif locations
Rank
Diffu_LP_319 (Barcode: GCACCTAA) TAGGAGG 1
Diffu_LP_387 (Barcode: CACGTACG) TAGGAGG 3
Diffu_LP_473 (Barcode: CTCTTAGA) TAGGAGG 4
Diffu_LP_472 (Barcode: GATTGACG) AAGGAGT 5
Diffu_LP_466 (Barcode: CAGCAACG) AAGGAGG 6
Diffu_LP_412 (Barcode: CGACATCA) AAGGTGA 8
Diffu_LP_70 (Barcode: AGCTAGGG) AAGGAGA 11
Diffu_LP_250 (Barcode: TTGGGCAC) AAAGAGG 14
Diffu_LP_47 (Barcode: GATCTTGA) TAGGAGA 17
Diffu_LP_329 (Barcode: TCCTATCA) AAGGAGA 18
Diffu_LP_77 (Barcode: ATTCACGA) AAGGAGA 20
Diffu_LP_82 (Barcode: CAAGGTAG) GGGGAGG 21
Diffu_LP_169 (Barcode: CTTACCAC) AAGGGGG 22
Diffu_LP_4 (Barcode: CGATCCTT) GAGGTGA 23
Diffu_LP_254 (Barcode: CACACGTC) GAGGGGT 24
Diffu_LP_474 (Barcode: TTACAGCC) AAGGTGA 25
d
Lactobacillus top promoters → B. thetaiotaomicron chassis
NanoLuc Luminescence (RLU/OD600)
0.0002
0.0012
262.32x
4.44x
1.37x
2.75x
1.29x
WT
P AM4-rplL*
P Diffu_LP_319
P Diffu_LP_235
P Diffu_LP_387
P Diffu_LP_473
P Diffu_LP_472

**Extended Data Fig. 8. Transcriptional activity and motif analysis of *Lactobacillus* promoters.** **a,** Distribution of transcriptional activity scores. **b,** Correlation of transcriptional activity between biological replicates. **c,** Motif locations within high-activity promoters. **d,** Reporter output of the top five *Lactobacillus* promoters in the heterologous host, *B. thetaiotaomicron*. Bars show mean ± SE; white circles represent individual biological replicates (n = 3 per group). Statistical significance was determined by an unpaired, two-sided Welch's t-test.

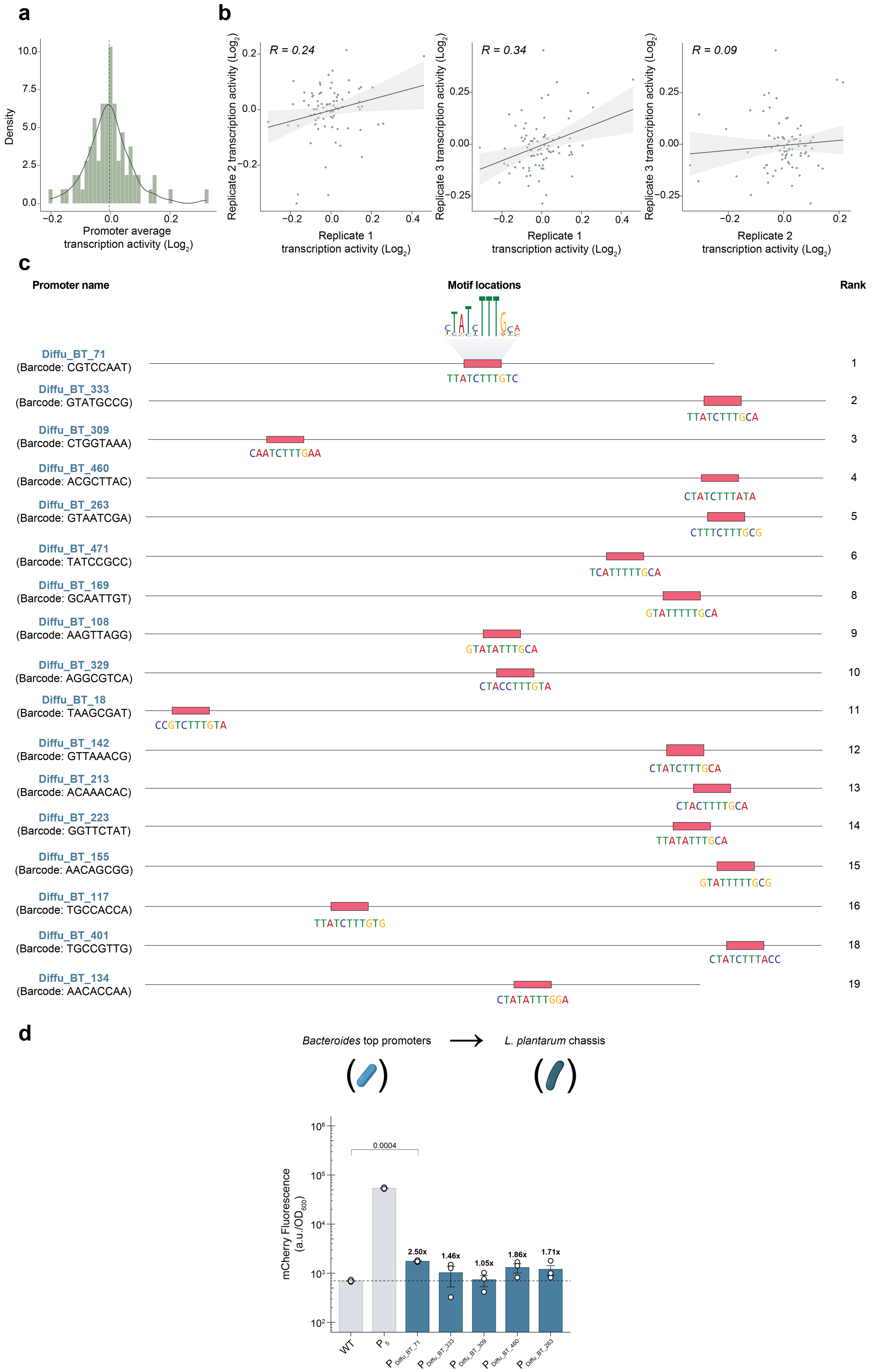
a
Density
10.0
7.5
5.0
2.5
0.0
−0.2
0.0
0.2
Promoter average
transcription activity (Log2)
b
R = 0.24
Replicate 2 transcription activity (Log2)
Replicate 1
transcription activity (Log2)
R = 0.34
Replicate 3 transcription activity (Log2)
Replicate 1
transcription activity (Log2)
R = 0.09
Replicate 3 transcription activity (Log2)
Replicate 2
transcription activity (Log2)
c
Promoter name
Motif locations
Rank
Diffu_BT_71
(Barcode: CGTCCAAT)
TTATCTTTGTC
1
Diffu_BT_333
(Barcode: GTATGCCG)
TTATCTTTGCA
2
Diffu_BT_309
(Barcode: CTGGTAAA)
CAATCTTTGAA
3
Diffu_BT_460
(Barcode: ACGCTTAC)
CTATCTTTATA
4
Diffu_BT_263
(Barcode: GTAATCGA)
CTTTCTTTGCG
5
Diffu_BT_471
(Barcode: TATCCGCC)
TCATTTTTGCA
6
Diffu_BT_169
(Barcode: GCAATTGT)
GTATTTTTGCA
8
Diffu_BT_108
(Barcode: AAGTTAGG)
GTATATTTGCA
9
Diffu_BT_329
(Barcode: AGGCGTCA)
CTACCTTTGTA
10
Diffu_BT_18
(Barcode: TAAGCGAT)
CCGTCTTTGTA
11
Diffu_BT_142
(Barcode: GTTAAACG)
CTATCTTTGCA
12
Diffu_BT_213
(Barcode: ACAAACAC)
CTACTTTTGCA
13
Diffu_BT_223
(Barcode: GGTTCTAT)
TTATATTTGCA
14
Diffu_BT_155
(Barcode: AACAGCGG)
GTATTTTTGCG
15
Diffu_BT_117
(Barcode: TGCCACCA)
TTATCTTTGTG
16
Diffu_BT_401
(Barcode: TGCCGTTG)
CTATCTTTACC
18
Diffu_BT_134
(Barcode: AACACCAA)
CTATATTTGGA
19
d
Bacteroides top promoters → L. plantarum chassis
mCherry Fluorescence
(a.u./OD600)
0.0004
2.50x
1.46x
1.05x
1.86x
1.71x
WT
P5
P Diffu_BT_71
P Diffu_BT_333
P Diffu_BT_309
P Diffu_BT_460
P Diffu_BT_263

**Extended Data Fig. 9. Transcriptional activity and motif analysis of *Bacteroides* promoters.** **a,** Distribution of transcriptional activity scores. **b,** Correlation of transcriptional activity between biological replicates. **c,** Motif locations within high-activity promoters. **d,** Reporter output of the top five *Bacteroides* promoters in the heterologous host, *L. plantarum*. Bars show mean ± SE; white circles represent individual biological replicates (n = 3 per group). Statistical significance was determined by an unpaired, two-sided Welch's t-test.

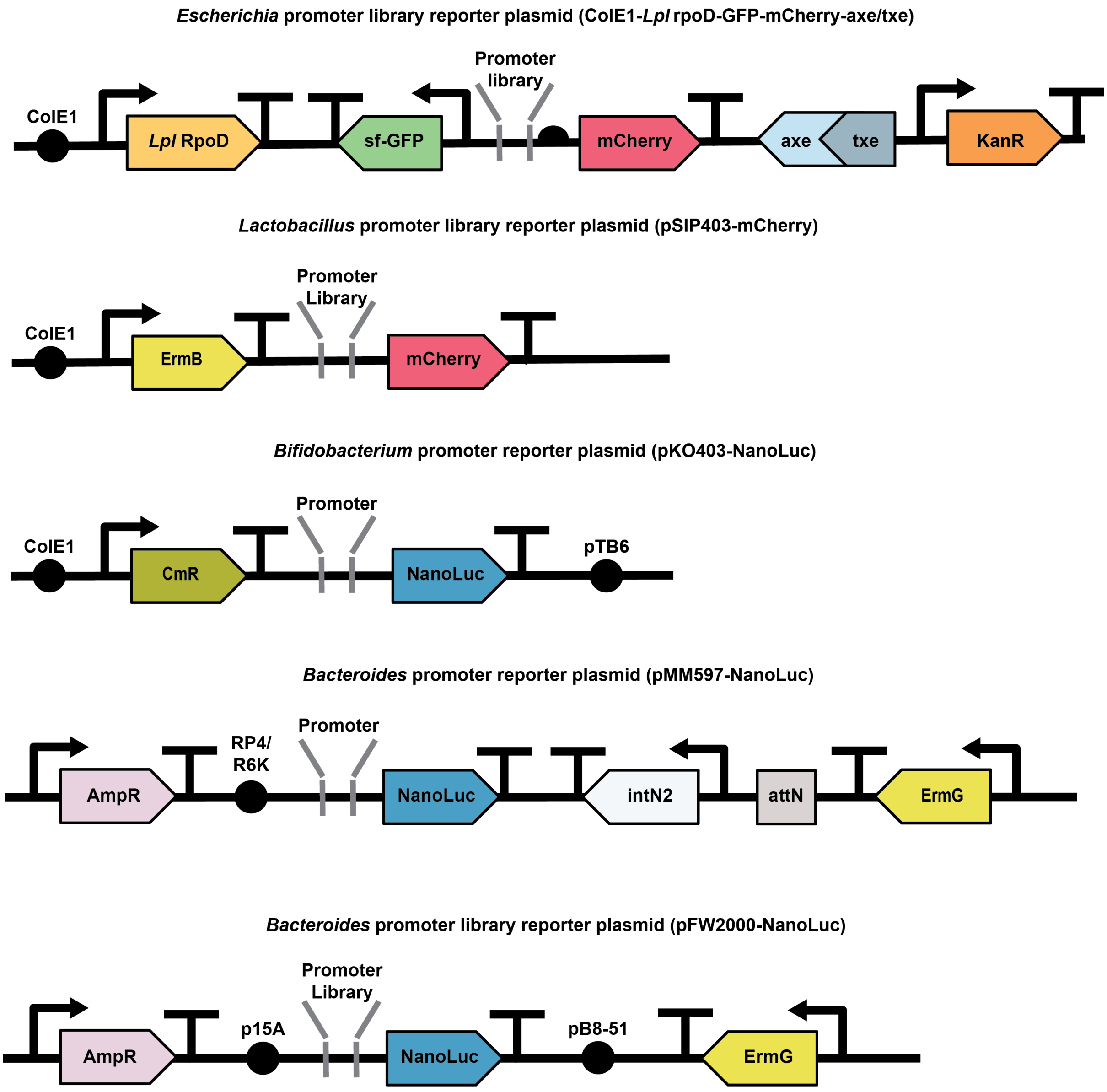


**Extended Data Fig. 10. Design of plasmid vectors used in this study.** Schematic maps of the expression constructs, including the promoter insertion sites, reporter genes, selection marker, and other components.
