## Supplementary Table 1 for "DAPE-PD: Diffusion adaptation of genomic foundation model enables tailored-generation of *cis*-regulatory DNA"

### Supplementary Table 1.
#### Bacterial Strains, Plasmids, Primers, and Genes Used in This Study

**Bacterial Strains Used in This Study**

| Species | Strain | Growth Conditions | Usages |
|---|---|---|---|
| *E. coli* | TOP10 | Aerobic, 37°C, 250 RPM | Preparation of chemically competent cells of molecular cloning procedures |
| | 10-Beta | Aerobic, 37°C, 250 RPM | Preparation of electrocompetent cells of molecular cloning procedures |
| | S17-1 λpir | Aerobic, 37°C, 250 RPM | Conjugation experiments with *B. thetaiotaomicron* |
| | Nissle 1917 | Aerobic, 37°C, 250 RPM | *Escherichia* promoter screening |
| *L. plantarum* | NC8 | Aerobic, 30°C, static | *Lactobacillus* promoter screening |
| *B. infantis* | S12 | Anaerobic, 37°C, static | *Bifidobacterium* promoter screening |
| *B. thetaiotaomicron* | VPI-5482 | Anaerobic, 37°C, static | *Bacteroides* promoter screening |

**Plasmids Used in This Study**

| Identifier | Plasmid | Relevant Features |
|---|---|---|
| megF088 | ColE1-*LpI* rpoD-GFP-PDiffu_EC_157-mCherry-axe/txe | mCherry expressed from PDiffu_EC_157 (*Escherichia* diffusion library promoter selected via FACS-seq), KanR, ColE1-LpI rpoD-GFP-axe/txe backbone |
| megF089 | ColE1-*LpI* rpoD-GFP-PDiffu_EC_462-mCherry-axe/txe | mCherry expressed from PDiffu_EC_462 (*Escherichia* diffusion library promoter selected via FACS-seq), KanR, ColE1-LpI rpoD-GFP-axe/txe backbone |
| megF090 | ColE1-*LpI* rpoD-GFP-PDiffu_EC_179-mCherry-axe/txe | mCherry expressed from PDiffu_EC_179 (*Escherichia* diffusion library promoter selected via FACS-seq), KanR, ColE1-LpI rpoD-GFP-axe/txe backbone |
| megF091 | ColE1-*LpI* rpoD-GFP-PProD-mCherry-axe/txe | mCherry expressed from PProD (*E. coli* native constitutive promoter), KanR, ColE1-LpI rpoD-GFP-axe/txe backbone |
| megF042 | pSIP403-PDiffu_LP_1-mCherry | mCherry expressed from PDiffu_LP_1 (*Lactobacillus* diffusion library promoter selected via filtering pipeline), ErmB, pSIP403 backbone |
| megF043 | pSIP403-PDiffu_LP_2-mCherry | mCherry expressed from PDiffu_LP_2 (*Lactobacillus* diffusion library promoter selected via filtering pipeline), ErmB, pSIP403 backbone |
| megF044 | pSIP403-PDiffu_LP_4-mCherry | mCherry expressed from PDiffu_LP_3 (*Lactobacillus* diffusion library promoter selected via filtering pipeline), ErmB, pSIP403 backbone |
| megF045 | pSIP403-PDiffu_LP_3-mCherry | mCherry expressed from PDiffu_LP_4 (*Lactobacillus* diffusion library promoter selected via filtering pipeline), ErmB, pSIP403 backbone |
| megF046 | pSIP403-PDiffu_LP_5-mCherry | mCherry expressed from PDiffu_LP_5 (*Lactobacillus* diffusion library promoter selected via filtering pipeline), ErmB, pSIP403 backbone |
| megF092 | pSIP403-P5-mCherry | mCherry expressed from P5 (*L. plantarum* native constitutive promoter), ErmB, pSIP403 backbone |
| megF093 | pSIP403-PDiffu_LP_319-mCherry | mCherry expressed from PDiffu_LP_319 (*Lactobacillus* diffusion library promoter selected via high-throughput characterization), ErmB, pSIP403 backbone |
| megF094 | pSIP403-PDiffu_LP_235-mCherry | mCherry expressed from PDiffu_LP_235 (*Lactobacillus* diffusion library promoter selected via high-throughput characterization), ErmB, pSIP403 backbone |
| megF095 | pSIP403-PDiffu_LP_387-mCherry | mCherry expressed from PDiffu_LP_387 (*Lactobacillus* diffusion library promoter selected via high-throughput characterization), ErmB, pSIP403 backbone |
| megF096 | pSIP403-PDiffu_LP_473-mCherry | mCherry expressed from PDiffu_LP_473 (*Lactobacillus* diffusion library promoter selected via high-throughput characterization), ErmB, pSIP403 backbone |

| | | |
|---|---|---|
| megF097 | pSIP403-PDiffu_LP_472-mCherry | mCherry expressed from PDiffu_LP_472 (*Lactobacillus* diffusion library promoter selected via high-throughput characterization), ErmB, pSIP403 backbone |
| megF098 | pSIP403-PDiffu_BT_71-mCherry | mCherry expressed from PDiffu_BT_71 (*Bacteroides* diffusion library promoter selected via high-throughput characterization), ErmB, pSIP403 backbone |
| megF099 | pSIP403-PDiffu_BT_333-mCherry | mCherry expressed from PDiffu_BT_333 (*Bacteroides* diffusion library promoter selected via high-throughput characterization), ErmB, pSIP403 backbone |
| megF100 | pSIP403-PDiffu_BT_309-mCherry | mCherry expressed from PDiffu_BT_309 (*Bacteroides* diffusion library promoter selected via high-throughput characterization), ErmB, pSIP403 backbone |
| megF101 | pSIP403-PDiffu_BT_460-mCherry | mCherry expressed from PDiffu_BT_460 (*Bacteroides* diffusion library promoter selected via high-throughput characterization), ErmB, pSIP403 backbone |
| megF102 | pSIP403-PDiffu_BT_263-mCherry | mCherry expressed from PDiffu_BT_263 (*Bacteroides* diffusion library promoter selected via high-throughput characterization), ErmB, pSIP403 backbone |
| megF058 | pMM597-PDiffu_BT_1-NanoLuc | NanoLuc expressed from PDiffu_BT_1 (*Bacteroides* diffusion library promoter selected via filtering pipeline), AmpR (For selection in *E.coli*), ErmG (For selection in *B. thetaiotaomicron*), pMM597 backbone |
| megF059 | pMM597-PDiffu_BT_2-NanoLuc | NanoLuc expressed from PDiffu_BT_2 (*Bacteroides* diffusion library promoter selected via filtering pipeline), AmpR (For selection in *E.coli*), ErmG (For selection in *B. thetaiotaomicron*), pMM597 backbone |
| megF060 | pMM597-PDiffu_BT_3-NanoLuc | NanoLuc expressed from PDiffu_BT_3 (*Bacteroides* diffusion library promoter selected via filtering pipeline), AmpR (For selection in *E.coli*), ErmG (For selection in *B. thetaiotaomicron*), pMM597 backbone |
| megF061 | pMM597-PDiffu_BT_4-NanoLuc | NanoLuc expressed from PDiffu_BT_4 (*Bacteroides* diffusion library promoter selected via filtering pipeline), AmpR (For selection in *E.coli*), ErmG (For selection in *B. thetaiotaomicron*), pMM597 backbone |
| megF062 | pMM597-PDiffu_BT_5-NanoLuc | NanoLuc expressed from PDiffu_BT_5 (*Bacteroides* diffusion library promoter selected via filtering pipeline), AmpR (For selection in *E.coli*), ErmG (For selection in *B. thetaiotaomicron*), pMM597 backbone |
| megF087 | pMM597-PAM4-rpiL*-NanoLuc | NanoLuc expressed from PAM4-rpiL* (*B. thetaiotaomicron* native constitutive promoter), AmpR (For selection in *E.coli*), ErmG (For selection in *B. thetaiotaomicron*), pMM597 backbone |

| | | |
|---|---|---|
| megF114 | pMM597-PDiffu_LP_319-NanoLuc | NanoLuc expressed from PDiffu_LP_319 (*Lactobacillus* diffusion library promoter selected via high-throughput characterization), AmpR (For selection in *E.coli*), ErmG (For selection in *B. thetaiotaomicron*), pMM597 backbone |
| megF115 | pMM597-PDiffu_LP_235-NanoLuc | NanoLuc expressed from PDiffu_LP_235 (*Lactobacillus* diffusion library promoter selected via high-throughput characterization), AmpR (For selection in *E.coli*), ErmG (For selection in *B. thetaiotaomicron*), pMM597 backbone |
| megF116 | pMM597-PDiffu_LP_387-NanoLuc | NanoLuc expressed from PDiffu_LP_387 (Lactobacillus diffusion library promoter selected via high-throughput characterization), AmpR (For selection in E.coli), ErmG (For selection in B. thetaiotaomicron), pMM597 backbone |
| megF117 | pMM597-PDiffu_LP_473-NanoLuc | NanoLuc expressed from PDiffu_LP_473 (*Lactobacillus* diffusion library promoter selected via high-throughput characterization), AmpR (For selection in *E.coli*), ErmG (For selection in *B. thetaiotaomicron*), pMM597 backbone |
| megF118 | pMM597-PDiffu_LP_472-NanoLuc | NanoLuc expressed from PDiffu_LP_472 (*Lactobacillus* diffusion library promoter selected via high-throughput characterization), AmpR (For selection in *E.coli*), ErmG (For selection in *B. thetaiotaomicron*), pMM597 backbone |
| megF119 | pMM597-PDiffu_BT_71-NanoLuc | NanoLuc expressed from PDiffu_BT_71 (*Bacteroides* diffusion library promoter selected via high-throughput characterization), AmpR (For selection in *E.coli*), ErmG (For selection in *B. thetaiotaomicron*), pMM597 backbone |
| megF120 | pMM597-PDiffu_BT_333-NanoLuc | NanoLuc expressed from PDiffu_BT_333 (*Bacteroides* diffusion library promoter selected via high-throughput characterization), AmpR (For selection in *E.coli*), ErmG (For selection in *B. thetaiotaomicron*), pMM597 backbone |
| megF121 | pMM597-PDiffu_BT_309-NanoLuc | NanoLuc expressed from PDiffu_BT_309 (*Bacteroides* diffusion library promoter selected via high-throughput characterization), AmpR (For selection in *E.coli*), ErmG (For selection in *B. thetaiotaomicron*), pMM597 backbone |
| megF122 | pMM597-PDiffu_BT_460-NanoLuc | NanoLuc expressed from PDiffu_BT_460 (*Bacteroides* diffusion library promoter selected via high-throughput characterization), AmpR (For selection in *E.coli*), ErmG (For selection in *B. thetaiotaomicron*), pMM597 backbone |

| | | |
|---|---|---|
| megF123 | pMM597-PDiffu_BT_263-NanoLuc | NanoLuc expressed from PDiffu_BT_263 (*Bacteroides* diffusion library promoter selected via high-throughput characterization), AmpR (For selection in *E.coli*), ErmG (For selection in *B. thetaiotaomicron*), pMM597 backbone |
| megF125 | pKO403-PDiffu_BL_1-NanoLuc | NanoLuc expressed from PDiffu_BL_1 (*Bifidobacterium* diffusion library promoter selected via filtering pipeline), CmR, pKO403 backbone |
| megF126 | pKO403-PDiffu_BL_2-NanoLuc | NanoLuc expressed from PDiffu_BL_2 (*Bifidobacterium* diffusion library promoter selected via filtering pipeline), CmR, pKO403 backbone |
| megF127 | pKO403-PDiffu_BL_3-NanoLuc | NanoLuc expressed from PDiffu_BL_3 (*Bifidobacterium* diffusion library promoter selected via filtering pipeline), CmR, pKO403 backbone |
| megF128 | pKO403-PDiffu_BL_4-NanoLuc | NanoLuc expressed from PDiffu_BL_4 (*Bifidobacterium* diffusion library promoter selected via filtering pipeline), CmR, pKO403 backbone |
| megF129 | pKO403-PDiffu_BL_5-NanoLuc | NanoLuc expressed from PDiffu_BL_5 (*Bifidobacterium* diffusion library promoter selected via filtering pipeline), CmR, pKO403 backbone |
| megF130 | pKO403-PS17_gap-NanoLuc | NanoLuc expressed from PS17_gap (*B. infantis* native constitutive promoter), CmR, pKO403 backbone |
| *Escherichia* Diffusion FACS-seq Promoter Library | ColE1-*Lpl* rpoD-GFP-[Library Insertion Site]-mCherry-axe/txe | mCherry expression under the control of each of the *Escherichia* diffusion library promoters, KanR, ColE1-Lpl rpoD-GFP-axe/txe backbone |
| *Escherichia* AR FACS-seq Promoter Library | ColE1-*Lpl* rpoD-GFP-[Library Insertion Site]-mCherry-axe/txe | mCherry expression under the control of each of the *Escherichia* AR library (Joint + Separate) promoters, KanR, ColE1-Lpl rpoD-GFP-axe/txe backbone |
| *Lactobacillus* High-Throughput Characterization Promoter Library | pSIP403-[Library Insertion Site]-mCherry | mCherry expression under the control of each of the *Lactobacillus* diffusion library promoters, ErmB, pSIP403 backbone |
| *Bacteroides* High-Throughput Characterization Promoter Library | pFW2000-[Library Insertion Site]-NanoLuc | NanoLuc expression under the control of each of the *Bacteroides* diffusion library promoters, AmpR (For selection in *E.coli*), ErmG (For selection in *B. thetaiotaomicron*), pFW2000 backbone |

**NGS Primers Used in This Study**

Note: The primer attachment site on the template is denoted by lower-case letters, whereas the flanking adaptor sequences are shown in upper-case

| Name | Sequence | Description | Notes |
|---|---|---|---|
| megD001 | AATGATACGGCGACCACCGAGATCTACACTCGCCTTGACACTCTTTCCCTACACGACGCTCTTCCGATCTTggtgttagctgtggaaacaccg | NGS primer mix for Escherichia library FACS-seq assessment-Forward | A primer mix containing 10 forward staggered primers and one reverse primer was used for NGS sample preparation. This primer mix targets the promoter sequence in each of the library constructs |
| megD002 | AATGATACGGCGACCACCGAGATCTACACATAGCGTCACACTCTTTCCCTACACGACGCTCTTCCGATCTATggtgttagctgtggaaacaccg | | |
| megD003 | AATGATACGGCGACCACCGAGATCTACACGAAGAAGTACACTCTTTCCCTACACGACGCTCTTCCGATCTGATggtgttagctgtggaaacaccg | | |
| megD004 | AATGATACGGCGACCACCGAGATCTACACATTCTAGGACACTCTTTCCCTACACGACGCTCTTCCGATCTCGATggtgttagctgtggaaacaccg | | |
| megD005 | AATGATACGGCGACCACCGAGATCTACACCGTTACCAACACTCTTTCCCTACACGACGCTCTTCCGATCTTCGATggtgttagctgtggaaacaccg | | |
| megD006 | AATGATACGGCGACCACCGAGATCTACACGTCTGATGACACTCTTTCCCTACACGACGCTCTTCCGATCTATCGATggtgttagctgtggaaacaccg | | |
| megD007 | AATGATACGGCGACCACCGAGATCTACACTTACGCACACACTCTTTCCCTACACGACGCTCTTCCGATCTGATCGATggtgttagctgtggaaacaccg | | |
| megD008 | AATGATACGGCGACCACCGAGATCTACACTTGAATAGACACTCTTTCCCTACACGACGCTCTTCCGATCTCGATCGATggtgttagctgtggaaacaccg | | |
| megD009 | AATGATACGGCGACCACCGAGATCTACACTCCTTGGTACACTCTTTCCCTACACGACGCTCTTCCGATCTTCGATCGATggtgttagctgtggaaacaccg | | |
| megD010 | AATGATACGGCGACCACCGAGATCTACACACAGGTATACACTCTTTCCCTACACGACGCTCTTCCGATCTATCGATCGATggtgttagctgtggaaacaccg | | |
| megD016 | CAAGCAGAAGACGGCATACGAGATATGAAGTAGTGACTGGAGTTCAGACGTGTGCTCTTCCGATCTaagcacatccggtgacagct | NGS primers for *Escherichia* library FACS-seq assessment-Reverse | |
| megD053 | AATGATACGGCGACCACCGAGATCTACACTCGCCTTGACACTCTTTCCCTACACGACGCTCTTCCGATCTTcgagcagtgaacgcatgtatcc | NGS primers for *Lactobacillus* library high-throughput assessment-Forward | This primer pair targets the barcode region located between the reporter mCherry gene and the terminator in each of the library constructs |
| megD054 | CAAGCAGAAGACGGCATACGAGATTCCGTGCGGTGACTGGAGTTCAGACGTGTGCTCTTCCGATCTcgaacccggggtaccgaatt | NGS primers for *Lactobacillus* library high-throughput assessment-Reverse | |

| | | | |
|---|---|---|---|
| megD081 | AATGATACGGCGACCACCGAGATCTACACTCGCCTTGACACTCTTTCCCTACACGACGCTCTTCCGATCTTcggtacgctggtgattgatggt | NGS primers for *Bacteroides* library high-throughput assessment-Forward | This primer pair targets the barcode region located between the reporter NanoLuc gene and the terminator in each of the library constructs |
| megD082 | CAAGCAGAAGACGGCATACGAGATTCCGTGCGGTGACTGGAGTTCAGACGTGTGCTCTTCCGATCTagcactccgataacgattagagtgc | NGS primers for *Bacteroides* library high-throughput assessment-Reverse | |

**Reference Promoter Used**

| Promoter | Sequence |
|---|---|
| PProD | CACAGCTAACACCACGTCGTCCCTATCTGCTGCCCTAGGTCTATGAGTGGTTGCTGG<br>ATAACTTTACGGGCATGCATAAGGCTCGTATAATATATTCAGGGAGACCACAACGGTT<br>TCCCTCTACAAATAATTTTGTTTAACTTT |
| P5 | AGATCTCGGCCGGGAGTTGTTGACACACAAAACCAGACATGGTATTATAATCTATAG<br>CGAGGAGAAGATTTGTT |
| PAM4-rpiL* | TGATCTGGAAGAAGCAATGAAAGCTGCTGTTAAGTCTCCGAATCAGGTATTGTTCCTG<br>ACAGGTGTATTCCCATCCGGTAAACGCGGATACTTTGCAGTTGATCTGACTCAGGAAT<br>AAATTATAAATTAAGGTAAGAAGATTGTAGGATAAGCTAATGAAATAGAAAAAGGAT<br>GCCGTCACACAACTTGTCGGCATTCTTTTTTGTTTTATTAGTTGAAAATATAGTGAAAA<br>AGTTGCCTAAATATGTATGTTAACAAATTATTTGTCGTAACTTTGCACTCCCTTTGCAA<br>CAGCATAGCTCAGCACAGCGCATTTTAAAATAAAATAAATTATTTATTTAATTAAACGA<br>AT |
| PS17_gap | GCGGAATGCCTCGCATCGAATCGCCGCAGGCTGTACAGACATATTTGTTAGCGTTAA<br>CGAAATATGGCCGTTTTATGCTCAAAGCAAGCGCGACACCGTTGCTCTAGTACAGAC<br>GGCGCATTACAGTAGACACTGTTGGTAAACAAAGGCCATAGCGCATCCATGCGCAAA<br>CGGTCTACCCTACAAAGGGAGAACAC |

**Reporter Genes Used in This Study**

| Reporter Gene | Species | Sequence |
|---|---|---|
| mCherry | *E. coli* | ATGGTGAGCAAGGGCGAAGAAGATAACATGGCCATCATCAAGGAGTTCATGCGCTTCA<br>AGGTGCACATGGAGGGCTCCGTGAACGGCCACGAGTTCGAGATCGAGGGCGAGGG<br>CGAGGGCCGCCCCTACGAGGGCACCCAGACCGCCAAGCTGAAGGTGACCAAGGG<br>TGGCCCCCTGCCCTTCGCCTGGGACATCCTGTCCCCTCAGTTCATGTACGGCTCCAA<br>GGCCTACGTGAAGCACCCCGCCGACATCCCCGACTACTTGAAGCTGTCCTTCCCCG<br>AGGGCTTCAAGTGGGAGCGCGTGATGAACTTCGAGGACGGCGGCGTGGTGACCGTG<br>ACCCAGGACTCCTCCCTGCAGGACGGCGAGTTCATCTACAAGGTGAAGCTGCGCGG<br>CACCAACTTCCCCTCCGACGGCCCCGTAATGCAGAAGAAGACCATGGGCTGGGAGG<br>CCTCCTCCGAGCGGATGTACCCCGAGGACGGCGCCCTGAAGGGCGAGATCAAGCA<br>GAGGCTGAAGCTGAAGGACGGCGGCCACTACGACGCTGAGGTCAAGACCACCTACA<br>AGGCCAAGAAGCCCGTGCAGCTGCCCGGCGCCTACAACGTCAACATCAAGTTGGAC<br>ATCACCTCCCACAACGAGGACTACACCATCGTGGAACAGTACGAACGCGCCGAGGG<br>CCGCCACTCCACCGGCGGCATGGACGAGCTGTACAAGTAA |
| GFP | | ATGAGCAAAGGAGAAGAACTTTTCACTGGAGTTGTCCCAATTCTTGTTGAATTAGATGGT<br>GATGTTAATGGGCACAAATTTTCTGTCCGTGGAGAGGGTGAAGGTGATGCTACAAACGG<br>AAAACTCACCCTTAAATTTATTTGCACTACTGGAAAACTACCTGTTCCGTGGCCAACACT<br>TGTCACTACTCTGACCTATGGTGTTCAATGCTTTTCCCGTTATCCGGATCACATGAAACG<br>GCATGACTTTTTCAAGAGTGCCATGCCCGAAGGTTATGTACAGGAACGCACTATATCTTT<br>CAAAGATGACGGGACCTACAAGACGCGTGCTGAAGTCAAGTTTGAAGGTGATACCCTT<br>GTTAATCGTATCGAGTTAAAGGGTATTGATTTTAAAGAAGATGGAAACATTCTTGGACACA<br>AACTCGAGTACAACTTTAACTCACACAATGTATACATCACGGCAGACAAACAAAAGAAT<br>GGAATCAAAGCTAACTTCAAAATTCGCCACAACGTTGAAGATGGTTCCGTTCAACTAGC<br>AGACCATTATCAACAAAATACTCCAATTGGCGATGGCCCTGTCCTTTTACCAGACAACC<br>ATTACCTGTCGACACAATCTGTCCTTTCGAAAGATCCCAACGAAAAGCGTGACCACAT<br>GGTCCTTCTTGAGTTTGTAACTGCTGCTGGGATTACACATGGCATGGATGAGCTCTACA<br>AAGGATCCCATGGTACGCGTGCTAGAGGCATCAAATAA |
| mCherry | *L. plantarum* | ATGAATTCAGATATGGTTTCAAAAGGGGAGGAGGATAATATGGCGATTATCAAGGAATTT<br>ATGCGGTTTAAAGTGCATATGGAAGGGTCGGTCAATGGTCATGAATTTGAGATTGAAGGT<br>GAGGGCGAAGGGCGTCCGTATGAAGGTACGCAGACGGCGAAGTTAAAAGTCACTAAG<br>GGAGGTCCGCTGCCATTTGCTTGGGATATTTTGAGTCCACAGTTCATGTATGGATCGAA<br>AGCTTACGTAAAGCACCCTGCCGATATTCCGGATTATTTGAAGTTAAGTTTTCCAGAAGG<br>ATTTAAGTGGGAACGTGTTATGAACTTTGAAGACGGTGGCGTTGTTACAGTAACTCAAGA<br>TTCTTCGTTGCAAGATGGTGAATTTATTTACAAGGTTAAGTTGCGAGGTACGAATTTTCCG<br>TCAGATGGACCAGTAATGCAAAAGAAAACTATGGGTTGGGAAGCGAGCAGTGAACGCA<br>TGTATCCAGAAGATGGCGCTTTGAAAGGTGAAATTAAACAGCGGTTGAAATTAAAAGATG<br>GTGGTCACTATGATGCGGAAGTCAAAACGACTTATAAAGCTAAGAAACCAGTTCAACTG<br>CCAGGCGCATATAACGTCAATATTAAGCTGGATATCACGAGTCATAATGAAGACTATAC<br>GATCGTCGAACAGTATGAACGAGCAGAGGGACGACACTCAACAGGTGGCATGGACG<br>AATTATATAAATGA |
| NanoLuc | *B. infantis* | ATGGTTTTCACTTTAGAAGATTTCGTTGGTGATTGGCGTCAAACTGCTGGTTACAACTTAG<br>ATCAAGTTTTAGAACAAGGTGGTGTTTCATCATTATTCCAAAACTTAGGTGTTTCAGTTACT<br>CCAATCCAACGTATCGTTTTATCAGGTGAAAACGGTTTAAAGATCGATATCCACGTTATC<br>ATCCCATACGAAGGTTTATCAGGTGATCAAATGGGTCAAATCGAAAAGATCTTCAAGGTT<br>GTTTACCCAGTTGATGATCACCACTTCAAGGTTATCTTACACTACGGTACTTTAGTTATCG<br>ATGGTGTTACTCCAAACATGATCGATTACTTCGGTCGTCCATACGAAGGTATCGCTGTTT<br>TCGATGGTAAGAAGATCACTGTTACTGGTACTTTATGGAACGGTAACAAGATCATCGATG<br>AACGTTTAATCAACCCAGATGGTTCATTATTATTCCGTGTTACTATCAACGGTGTTACTGG<br>TTGGCGTTTATGTGAACGTATCTTAGCTTAA |

| | | |
|---|---|---|
| NanoLuc | *B. thetaiotaomicron* | ATGGTTTTTACTCTGGAAGATTTTGTTGGCGATTGGCGTCAGACCGCGGGTTATAATTTG<br>GATCAAGTCCTGGAACAGGGTGGCGTAAGCTCTCTGTTCCAGAACCTGGGTGTGAGC<br>GTGACGCCGATTCAGCGCATCGTTCTGTCCGGCGAGAACGGTCTGAAAATTGATATTC<br>ATGTGATCATCCCGTACGAAGGCCTGAGCGGTGACCAAATGGGTCAAATCGAGAAAAT<br>CTTTAAAGTCGTCTACCCAGTTGACGATCACCACTTCAAGGTTATCTTGCATTACGGTAC<br>GCTGGTGATTGATGGTGTGACCCCGAATATGATTGACTATTTCGGCCGTCCGTATGAAG<br>GCATTGCCGTTTTTGACGGTAAAAAGATCACCGTCACCGGTACCCTGTGGAATGGCAA<br>TAAGATTATTGACGAGCGTCTGATTAACCCGGACGGCAGCCTGCTGTTCCGCGTGACC<br>ATCAACGGTGTCACGGGTTGGCGTCTGTGCGAGCGCATCCTGGCATAA |
